# Colocalization of Cancer Associated Biomarkers on Single Extracellular Vesicles for Early Cancer Detection

**DOI:** 10.1101/2023.02.07.527360

**Authors:** Daniel P. Salem, Laura T. Bortolin, Dan Gusenleitner, Jonian Grosha, Ibukunoluwapo O. Zabroski, Kelly M. Biette, Sanchari Banerjee, Christopher R. Sedlak, Delaney M. Byrne, Bilal F. Hamzeh, MacKenzie S. King, Lauren T. Cuoco, Timothy Santos-Heiman, Peter A. Duff, Emily S. Winn-Deen, Toumy Guettouche, Dawn R. Mattoon, Eric K. Huang, Randy Schekman, Anthony D. Couvillon, Joseph C. Sedlak

**Author notes:** co-first authors (Daniel P. Salem, Laura T. Bortolin).

## Abstract

Detection of cancer early, when it is most treatable, remains a significant challenge due to the lack of diagnostic methods sufficiently sensitive to detect nascent tumors. Early-stage tumors are small relative to their tissue of origin, heterogeneous, and infrequently manifest in clinical symptoms. Detection of their presence is made more difficult by a lack of abundant tumor-specific indicators (i.e., protein biomarkers, circulating tumor DNA, etc.) that would enable detection using a non-invasive diagnostic assay. In addition, many benign conditions manifest in a similar manner, thus discriminating an early-stage cancerous lesion from a benign tumor can present additional challenges and result in unnecessary medical procedures. To overcome these obstacles, we have developed a liquid biopsy assay that interrogates circulating extracellular vesicles (EVs) to detect tumor-specific biomarkers colocalized on the surface of individual EVs. Extracellular vesicles from all cell types, including early-stage tumors, are known to be abundant in blood, are remarkably stable, and serve as a biopsy of their cell of origin. The detection of a colocalized combination of cancer associated biomarkers that provide tumor specificity on the surface of extracellular vesicles enables the discrimination of early- and late-stage cancer from non-malignant conditions.

## Introduction

Cancer remains a significant public health challenge globally and is the second leading cause of death in the United States after heart disease (Siegel et al., 2023). Significant advances in the availability and efficacy of therapy have improved survival for some cancers, but the economic burden associated with cancer care has risen precipitously, and is projected to approach $245 billion by 2030, with annual and cumulative costs of care increasing with cancer stage (Mariotto et al., 2020; McGarvey et al., 2022). Cancer diagnosed at an early stage is addressable through an array of treatment modalities and is typically more responsive to therapy (Siegel et al., 2023; Sung et al., 2021). Given the significant clinical and economic benefits, the development and implementation of improved diagnostic tools for the early detection of cancer is a public health imperative.

To deliver net clinical benefit at a population scale, cancer screening tests must incorporate a foundational set of features. Tests intended for the early detection of cancer must have sufficient sensitivity to identify small, early-stage tumors, and sufficient specificity to minimize over-diagnosis and false positives due to confounding co-morbidities that may be present in the target population. Tests must be designed to facilitate broad adoption, including geographic and economic accessibility, and an experience that is convenient and noninvasive for the patient. Ideally, diagnostic tests also offer insights into the molecular mechanisms of the underlying cancer that can help guide clinical management. Tests designated for screening in otherwise healthy asymptomatic individuals, whether deployed in an average-risk or high-risk patient population, have the added requirement of demonstrated mortality benefit, requiring large, randomized control trials (Crosby et al., 2022). To date, such trials have only been successfully completed for select high-prevalence cancers, resulting in routine screening recommendations only for colorectal, lung, and breast cancers (US Preventive Services Task Force, 2009; US Preventive Services Task Force et al., 2021b, 2021a). As a result, more than 70% of cancer deaths in the United States result from malignancies for which there is no recommendation for screening (Beer, 2021).

Traditional diagnostic modalities aren’t well suited for early cancer detection. Imaging-based techniques require specialized equipment that may not be broadly accessible, and often necessitate a separate clinical appointment. Low-cost imaging methods such as low-dose computed tomography (LDCT) for lung cancer screening, and trans-vaginal ultrasound for ovarian cancer screening lack the specificity to accurately differentiate malignancies from benign masses (Pavlik and van Nagell, 2013; Pinsky, 2014; van Nagell et al., 2007). While higher resolution imaging techniques such as PET/CT help to address this lack of specificity, the higher cost and more limited availability make them poorly suited as front-line diagnostic tools. Blood-based biomarker tests are an attractive alternative to imaging due to the low cost and broad accessibility, but single biomarker tests including PSA for prostate cancer screening and CA125 for ovarian cancer screening have exhibited limited specificity and their use has not been associated with a demonstrated mortality benefit (Menon et al., 2021, 2009; US Preventive Services Task Force et al., 2018). More recently, liquid biopsy approaches have been developed that are designed to measure genomic material released into circulation by apoptotic tumor cells. These methods leverage next generation sequencing (NGS) to measure a variety of genomic features including the presence of somatic mutations, methylation patterns, and fragment lengths, in some cases integrating thousands of features into a classifier that yields a determination of the presence or absence of cancer (Ignatiadis et al., 2021; Lone et al., 2022). In addition to the cost and complexity associated with NGS-based testing, these methods must overcome the biological limitation associated with the low abundance of circulating tumor DNA present in the context of small, early-stage tumors, and the limited stability of genomic material in standard blood collection tubes. These challenges may be addressed through the testing of large volumes of blood collected in specialized collection tubes designed to stabilize circulating DNA and RNA but introduce new logistic considerations (Dang and Park, 2022; Desai and Lovly, 2023; Yu et al., 2022, 2021). Finally, classifiers trained on thousands of features run the risk of overfitting to chance effects, and even if proper statistical care is taken, the detected effects may be statistically valid but biologically false (e.g., predicting slight differences in sample storage or handling instead of the disease, or other confounders). While these challenges can be overcome with large sample sizes and rigorous validation protocols, it makes such classifiers difficult to develop, validate and operationalize in practice (Cabús et al., 2022; Han and Jiang, 2014).

Extracellular vesicles (EVs) represent a diverse class of lipid-bilayer bounded nanoparticles ranging from 30 to 10,000 nanometers in diameter (van Niel et al., 2018; Willms et al., 2018). They are released by all living cells and serve an array of biological functions including cell-to-cell communication and the facilitation of cancer progression by establishing a metastatic niche and suppressing the immune response (Bebelman et al., 2018; Jerabkova-Roda et al., 2022; Lone et al., 2022; van Niel et al., 2018; Willms et al., 2018). EVs have been shown to contain numerous biomarkers derived from their cell of origin including proteins, lipids, and nucleic acids (Ghodasara et al., 2023). Tumor derived EVs are a promising analyte for early cancer detection due to their abundance in blood and molecular similarity with their cell of origin. Unlike genomic-based detection methods, cell death is not required for the release of EVs. (Amelio al., 2020; Yu et al., 2022) As a result, tumor-derived EVs outnumber tumor-derived DNA copies in circulation by orders of magnitude in early-stage disease (Avanzini et al., 2020; Ferguson and Weissleder, 2020). This enables the utilization of smaller sample volumes (∼100 µL) relative to genomic-based liquid biopsy assays, which require several milliliters of blood per run (Liu et al., 2021). The high plasma EV concentration of ∼10^10^ EVs per mL and estimated tumor-derived EV shedding rates per cubic millimeter of tumor volume make EVs an abundant source of tumor-derived biomarkers to target in cancer screening assays designed for detection of smaller, early-stage tumors (Ferguson et al., 2022; Ferguson and Weissleder, 2020; Johnsen et al., 2019). Given their abundance, stability, and molecular similarity to their cell of origin, EVs offer a unique diagnostic test analyte that may offer advantages over other modalities.

We have developed a method that interrogates the billions of EVs within a plasma or serum sample for the simultaneous detection of colocalized, cancer-associated biomarkers on the surface of individual EVs. Following the rational design of biomarker combinations, the clinical feasibility of this approach was applied to the detection of high-grade serous ovarian cancer (HGSOC), the most prevalent and deadliest histotype of ovarian cancer (Koshiyama et al., 2017).

## Results

### Measurement of colocalized biomarkers using immunocapture and detection

A conceptual schematic of the assay is depicted in Figure 1. EVs from cultured human cell lines, human plasma, or serum are enriched using size exclusion chromatography (SEC, Figure 1A). For capture and detection of biomarkers on the surface of EVs, monoclonal antibodies specific to the extracellular domains of cancer-associated targets are conjugated to magnetic beads (capture antibody) or complementary dsDNA oligonucleotides (detection antibody) as indicated in Figure 1B. Enriched EVs are isolated using a antibodies to the capture biomarker (Figure 1C) followed by detection with two antibodies to a second and third biomarker conjugated to complementary dsDNA oligonucleotides (Figure 1D). These complementary oligonucleotides hybridize and ligate if two DNA-conjugated antibodies are bound in proximity on the same EV (Figure 1D). The ligated DNA contains primer sites serving as a template for qPCR amplification to quantify the number of colocalization events in the sample. Based on this assay design, the complete set of biomarkers, detected by the antibody conjugated to the magnetic bead as well as the antibodies conjugated to the complementary oligonucleotides (a “biomarker combination”)—must be present on the surface of a single EV to generate a signal. In the following figures the assay readout is generally presented as a boxplot using the PCR cycle threshold (Ct) to discriminate positive signal (low Ct) from negative signal (high Ct).

**Figure 1.**
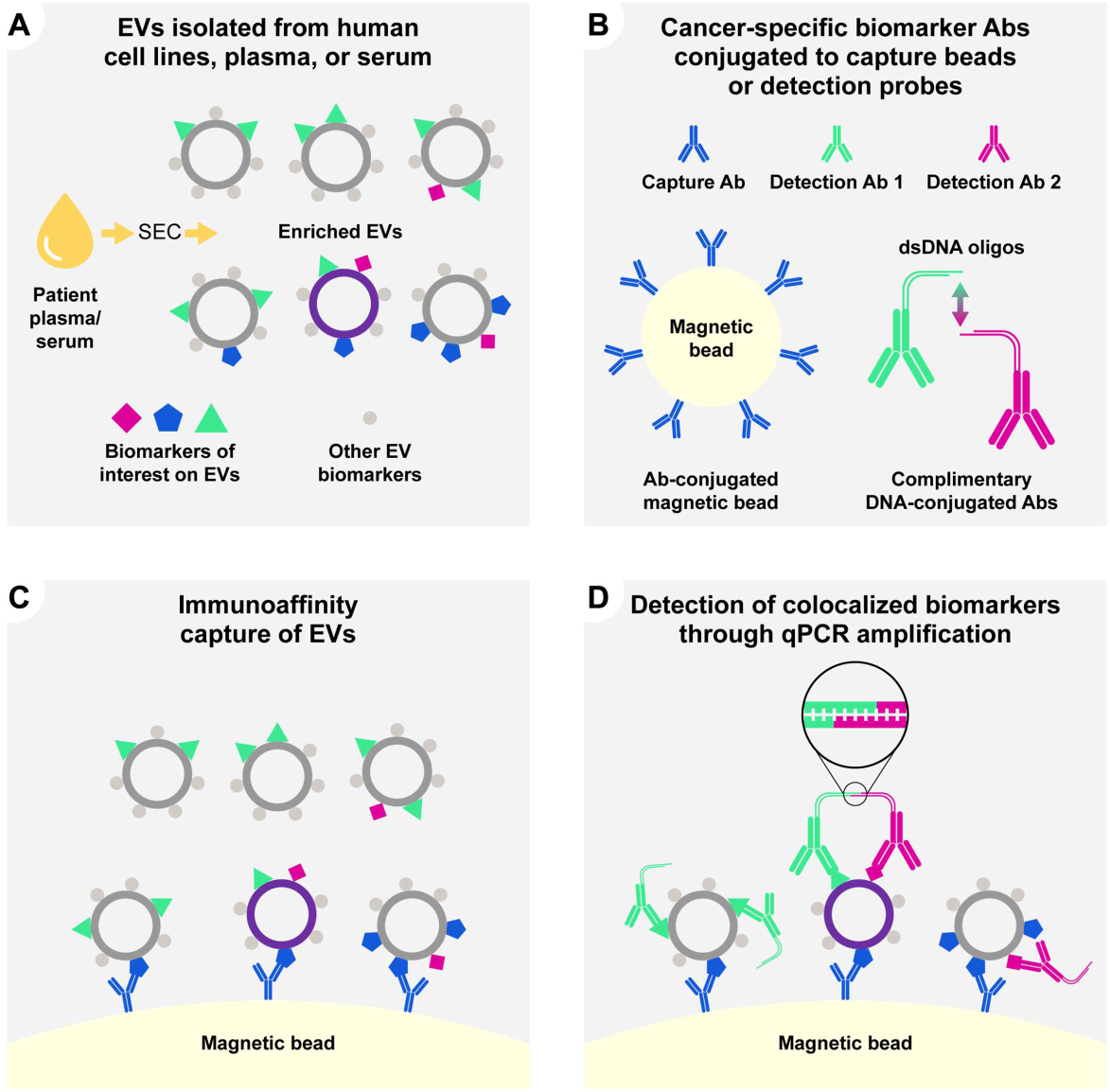
Overview of the assay method. Schematic representation of the assay workflow. (A) Extracellular vesicles are enriched from human cell line conditioned media, plasma, or serum by size-exclusion chromatography (SEC). Enriched EVs contain cargo and surface biomarkers from their cell-of-origin. (B) Antibodies targeting cancer-associated biomarker antibodies are conjugated to magnetic beads (capture antibody) or dsDNA oligonucleotides (detection antibodies). (C) SEC-enriched EVs are captured in solution with magnetic bead-antibody conjugates targeting a specific surface biomarker. (D) Immunoaffinity captured EVs are incubated with detection antibodies conjugated to complementary double-stranded DNA probes. The dsDNA oligonucleotides contain single-stranded overhangs which ligate only when in proximity to a complementary probe to generate a template for PCR. The abundance of the detection biomarkers captured on the EVs is read out using quantitative PCR. Positive signal or low cycle threshold (low Ct) is the result of all three biomarker antibody epitopes being present simultaneously on the surface of the same EV, the absence of one or more biomarkers results in failed capture or failed detection and low assay signal (high Ct).

### Identification and selection of biomarkers

Bone marrow stromal antigen-2/CD317/Tetherin (BST2), Folate receptor alpha (FOLR1), and Mucin-1 (MUC1) are cell surface proteins previously shown to be overexpressed in ovarian cancer (Gautam et al., 2023; Giampaolino et al., 2020, 2019; Januchowski et al., 2017; Kufe, 2009; Trinidad et al., 2023; Varaganti et al., 2023; Yang et al., 2022). Protein expression of all three biomarkers was demonstrated by western blotting to be present in the EV fractions from human ovarian cancer cell lines (Figure 2) and human plasma (not shown) enriched by size exclusion.

**Figure 2.**
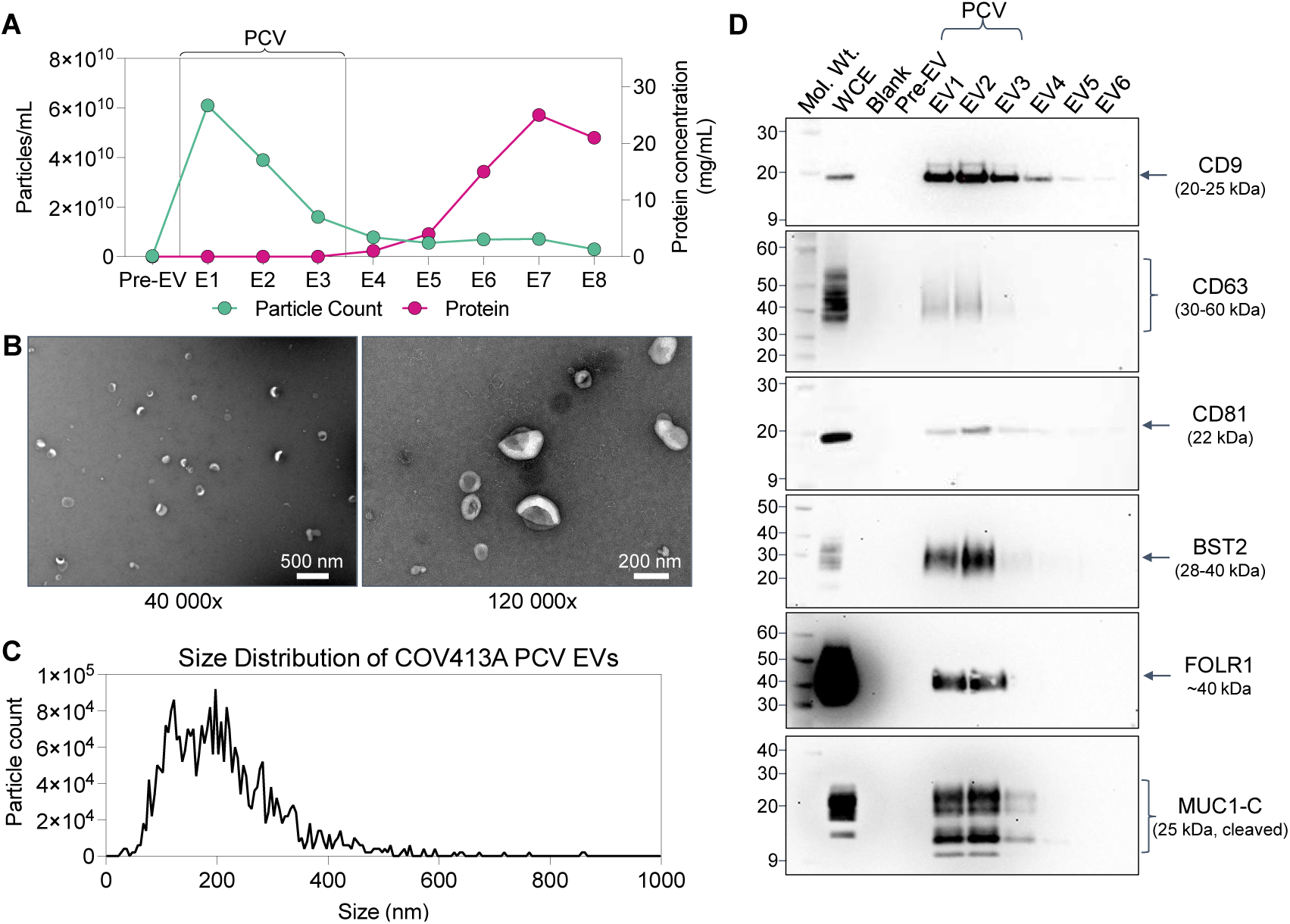
Characterization of human cancer cell line extracellular vesicles. SEC-enriched EV fractions from human ovarian cancer cell line (COV413A) conditioned media were characterized according to MISEV guidelines. (A) Nanoparticle tracking analysis (NTA, grey bars) and protein quantification (black line) of each fraction collected from the SEC column demonstrating the bulk of the nanoparticles elute in EV1-EV3, distinct from the bulk of soluble protein (EV6-EV8). The first fraction (Pre-EV) is the sum of the void and wash volumes of the column which generally contains larger particles, such as lipoproteins, which are not observed in cell line conditioned media. The purified collection volume (PCV), fractions EV1-EV3, containing the EVs used in downstream assay steps are highlighted. (B) Transmission electron microscopy of EVs derived from the total PCV (pooled fractions EV1-EV3) from COV413A conditioned media at two different magnifications demonstrates intact EVs ranging from ∼40-300 nm. Scale bars are 500 nm (40,000x, left) and 200 nm (120,000x, right), respectively. (C) Particle size distribution of pooled SEC fractions EV1-EV3 demonstrating particles consistent with the size range of small EVs ranging from ∼40-400 nm. (D) Western blot analysis of each of the initial fractions from the SEC column (Pre-EV, EV1-EV6) from COV413A conditioned media demonstrating the presence of known EV surface biomarkers (CD9, CD63, CD81), as well as BST2, FOLR1, and MUC1. Note that an antibody detecting the cleaved fragment of MUC1 (MUC1-C) was used to facilitate immunoblotting due to the very high molecular weight of MUC1 (250-500 kDa). Evaluation of additional MISEV protein markers, the presence of EV cargo proteins (flotillin, GAPDH) and absence of non-EV biomarkers (RPS3, Tom20) can be found in supplemental figure 2. Whole cell extracts (WCE, 10 ug total protein per lane) from COV413A were analyzed in parallel as a control for protein expression and biotinylated molecular weight (Mol. Wt.) markers were used to determine apparent molecular mass.

### Analyte Characterization

EVs enriched by SEC from human cell line conditioned media (Figure 2A, Supplemental Figure 2) and human plasma (Supplemental Figure 1) were characterized according to MISEV2018 guidelines by nanoparticle tracking analysis (NTA), transmission electron microscopy (TEM), and western blot to evaluate particle size, number, morphology, and phenotype (Théry et al., 2018). NTA for EVs from human cell lines (Figure 2A, C) and human plasma (Supplemental Figure 2A and 2B) demonstrates EV SEC fractions 1-3 (EV1-3) contain particles ranging in size from 40-400 nm consistent with the known size distribution for EVs. We selected and pooled EV1-3 for immunocapture and detection in our assay. In human plasma, EV4 was excluded due to the higher background introduced by the abundance of soluble protein in human plasma (Supplemental Figure 2A) and absence of EV biomarkers (CD9, CD81) as determined by western blot (Supplemental Figure 2C). The fundamental differences in NTA counts demonstrated between human plasma (Supplemental Figure 2) and human cell line conditioned media (Figure 2) is explained by the difference in heterogeneity between nanoparticles derived from human blood versus those enriched from a clonal cell line in culture (Lobb et al., 2015; Veerman et al., 2021). Enriched nanoparticles from both cell line and plasma EV fractions were further characterized by TEM demonstrating lipid-bounded nanoparticles ranging from 40-400 nm consistent with the NTA results (Figure 2B and Supplemental Figure 2D)(Cizmar and Yuana, 2017; Veerman et al., 2021). Additionally, western blotting was used to confirm the presence of accepted EV markers and absence of non-EV proteins as well as the presence of BST2, FOLR1 and MUC1 proteins within the EV fraction (Figure 2D and Supplemental Figure 2).

### Antibody Screening

Commercially available monoclonal antibodies to FOLR1, BST2 and MUC1 were screened for their ability to capture or detect EVs enriched from human cell lines predicted to express high (positive cell line) or low/no (negative cell line) levels of the selected biomarker (Supplemental Figure 3). Representative screening data for anti-FOLR1 antibodies used for capture (Supplemental Figure 3A) and detection (Supplemental Figure 3B) are shown using EVs derived from COV413A (FOLR1 positive) and SK-MEL-1 (FOLR1 negative) cell lines. Of the six antibodies screened for capture using CD81 antibodies for detection, antibodies 1 and 5 were selected for additional characterization based on the signal to background difference between the positive and negative cell line among other criteria. Antibodies 4 and 6 illustrate examples of reagents that failed validation in the capture assay due to lack of signal in the positive cell line (Supplemental Figure 3A). Antibodies 1 and 5 were further validated for detection using anti-CD81 conjugated beads for capture prior to detection with anti-FOLR1 antibodies (Supplemental Figure 3B). As with capture performance, both antibodies perform well in the detection step as illustrated by their ability to discriminate FOLR1 positive cell line EVs (COV413A) from FOLR1 negative cell line EVs (SK-MEL-1). Screening data for antibodies against BST2 and MUC1 described in this study are shown in Supplemental Figure 3 C-F.

### Cancer biomarkers are colocalized on the surface of EVs

To demonstrate colocalization of cancer-associated biomarkers with canonical EV biomarkers, the assay was applied to EVs derived from human cell lines known to express high or low levels of each cancer-associated biomarker. After capturing SEC-enriched EVs with antibodies directed against canonical EV tetraspanin biomarkers (CD63 or CD81), bound EVs were detected using detection antibodies directed against the extracellular domains of BST2, FOLR1, or MUC1 (Figure 3A, B and C, respectively). Only EVs derived from cell lines positive for both the tetraspanin and the cancer-associated biomarker result in a strong qPCR signal.

**Figure 3:**
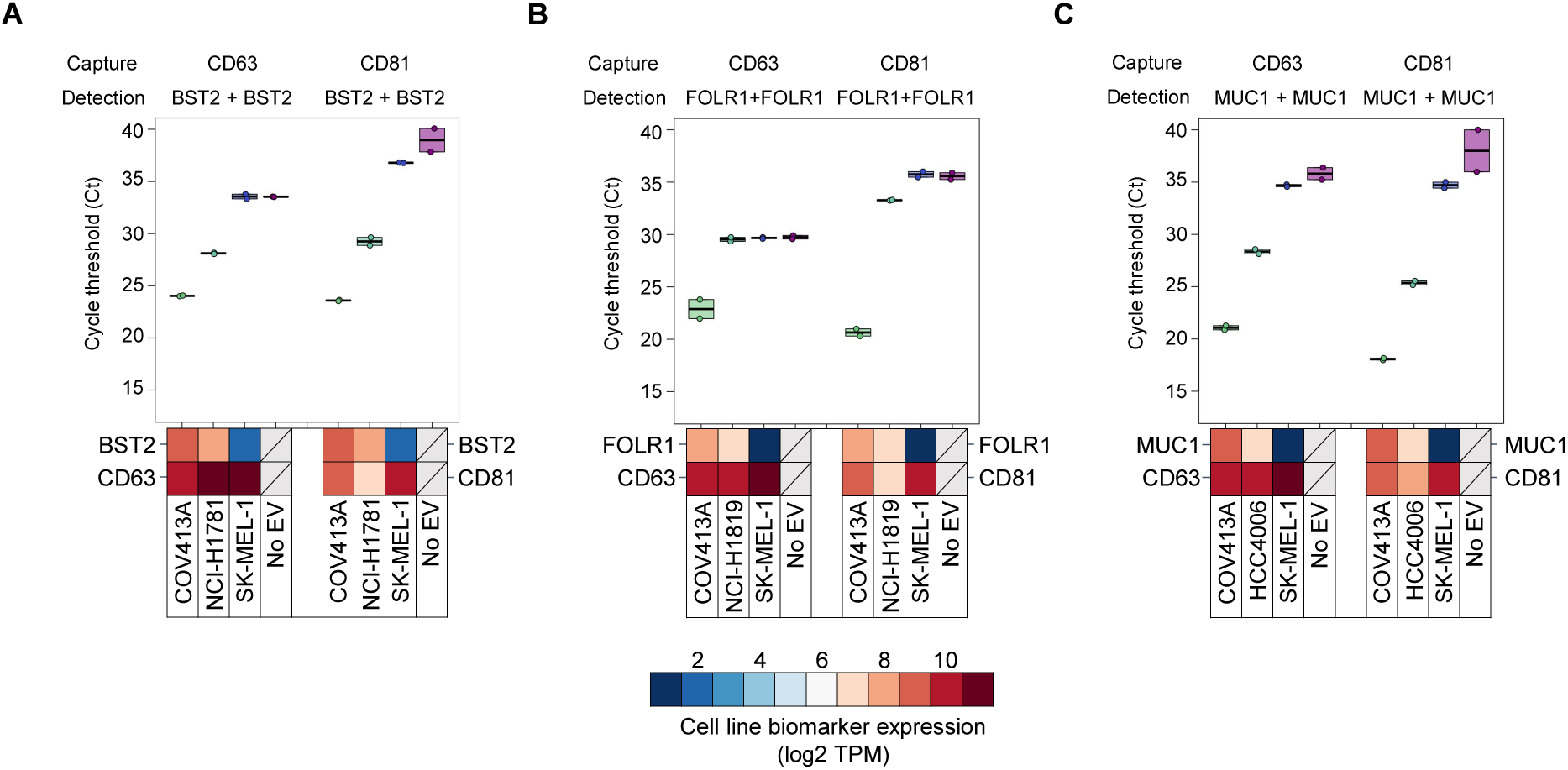
BST2, FOLR1, and MUC1 are colocalized with CD63 and CD81 in cell line EVs. EVs derived from cell lines predicted to express high (red), medium (white) or low (blue) levels of the indicated biomarkers based on mRNA analysis were captured with antibody-bead conjugates using anti-CD63 or anti-CD81 antibodies. Detection of bound EVs was determined using oligo-conjugated antibodies against (A) BST2, (B) FOLR1, or (C) MUC1. COV413A EVs were used as a positive control since this line expresses high levels of all three biomarkers studied. For BST2, FOLR1, and MUC1, a second positive cell line, NCI-H1781, NCI-H1819, and HCC-4006, respectively, was also included as a control. Note that despite moderate FOLR1 mRNA expression in NCI-H1819 cells, proteomics and western blot data would suggest otherwise (data not shown) suggesting this cell line may not express FOLR1 protein. SK-MEL-1, which expresses low levels of each marker was used as a negative control. Heat maps were included to demonstrate relative expression levels (log2 TPM) for each marker, expression data are derived from the CCLE database at the Broad Institute. No EV controls were used to eliminate antibodies incompatible with the assay (antibodies that demonstrated non-specific binding to the beads or other antibodies under the assay conditions).

Assay signal operates akin to a Boolean logic “AND gate” whereby all three surface epitopes must be confined to the same EV for PCR template generation to occur. To demonstrate this, we characterized EVs derived from four human cancer cell lines (COV413A, OVKATE, SW900, and G-401) exhibiting a range of BST2, FOLR1, and MUC1 expression levels (Figure 4A-C). We tested the functionality of the assay using two-biomarker (BST2 capture/FOLR1 + FOLR1 detection and BST2 capture/MUC1 + MUC1 detection) and three-biomarker (BST2 capture/FOLR1 + MUC1 detection) combinations in EVs from cell lines that were BST2^high^/FOLR1^high^/MUC1^low^ (OVKATE), BST2^high^/FOLR1^low^/MUC1^high^ (G-401 and SW900), and BST2^high^/FOLR1^high^/MUC1^high^ (COV413A) to demonstrate that assay signal is dependent on all of the capture and detection biomarkers being present for signal. Cell line EVs were tested at an equivalent concentration of 1.6×10^9^ particles/mL. As shown in Figure 4, signal was observed to correlate with biomarker colocalization in expected patterns. BST2 capture/FOLR1 + FOLR1 detection and BST2 capture/MUC1 + MUC1 detection generated strong signals for the cell lines that express both biomarkers (COV413A/OVKATE and COV413A/SW900, respectively). Weak BST2 capture/FOLR1 + FOLR1 detection and BST2 capture/MUC1 + MUC1 detection signals were observed for cell lines expressing low levels of at least one marker (SW900/G-401 and OVKATE/G-401, respectively). The three-biomarker combination BST2/FOLR1/MUC1 showed a clear separation between COV413A EVs (triple-positive) and cell lines missing at least one biomarker, suggesting assay signal is only robust when all biomarkers are colocalized on the same EV.

**Figure 4:**
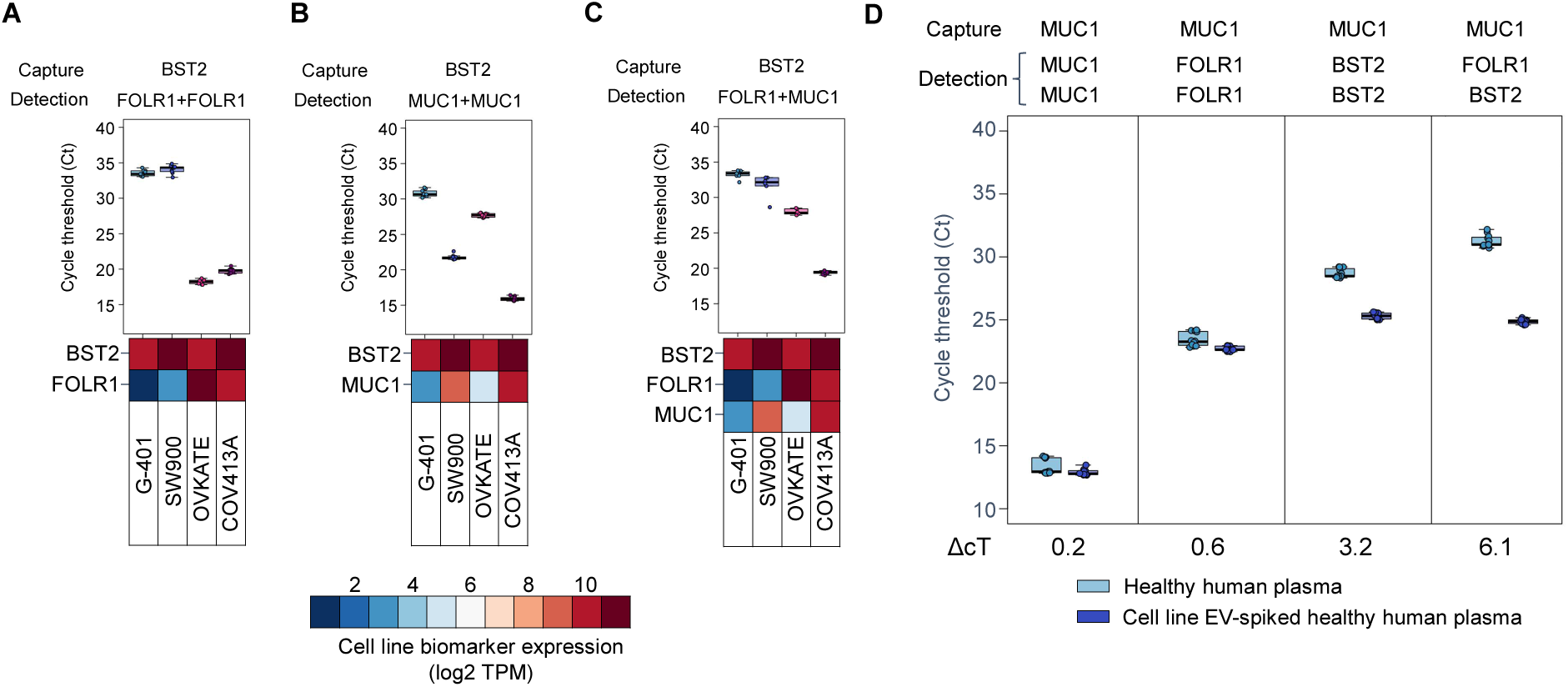
Colocalization of biomarkers is required for assay performance and improves signal to noise ratio. All three biomarkers must be present on the same EV for productive readout. If only two biomarkers are present on the EVs (one capture + one detect), no ligation occurs, and Ct is high. (A, B) Doublet (i.e. BST2 capture, FOLR1 + FOLR1 detection or BST2 capture, MUC1 + MUC1 detection) or (C) triplet combinations (BST2 capture, FOLR1 + MUC1 detection) of antibodies tested on cell line EVs from COV413A (BST2^high^/FOLR1^high^/MUC1^high^), OVKATE (BST2^high^/FOLR1^high^/MUC1^low^), SW900 (BST2^high^/FOLR1^low^/MUC1^high^), and G-401 (BST2^high^/FOLR1^low^/MUC1^low^) showing that doublets of BST2 capture/FOLR1 + FOLR1 detection (A), BST2 capture/MUC1 + MUC1 detection (B), or a triplet of BST2 capture/FOLR1 + MUC1 detection (C) only yield positive qPCR signal in cell lines with matching high biomarker expression. Biomarker mRNA abundance (log2 TPM) is represented for each biomarker in each cell line. (D) Signal from cell line EV-spiked (dark blue) or unspiked (light blue) healthy human plasma from a representative set of one-biomarker (MUC1 capture, MUC1 + MUC1 detection), two-biomarker (MUC1 capture/FOLR1 + FOLR1 detection or MUC1 capture/BST2 + BST2 detection), and three-biomarker (MUC1 capture/FOLR1 + BST2 detection) combinations. Increasing the number of unique, colocalized biomarkers results in reduced healthy background and improved discrimination of cancer cell line EV-spiked plasma. Signal difference (ΔCt) between unspiked healthy human plasma and spiked plasma is indicated.

### Detecting multiple colocalized biomarkers on EVs improves the signal-to-noise ratio

We hypothesized that the colocalization of multiple cancer-associated biomarkers could improve the specificity of distinguishing dilute cancer EVs from normal EVs. To test this, we spiked plasma from healthy control subjects with EVs derived from ovarian cancer cell lines and evaluated discrimination performance using combinations of one, two, and three unique biomarkers. Spiked plasma was prepared by mixing purified COV413A EVs with a female plasma pool derived from ninety healthy women at a final concentration of 1×10^8^ COV413A particles/mL. Unspiked and spiked plasma pools were characterized in the assay using biomarker combinations containing BST2, FOLR1, and/or MUC1 in various capture/detection assay configurations. Single-biomarker combinations comprising capture and detection antibodies targeting MUC1 only (MUC1 capture, MUC1 + MUC1 detection, Figure 4D; left panel) BST2-only, or FOLR1-only (data not shown), generally exhibited poor discrimination between unspiked and spiked plasma samples (median ΔCt = 0.2). However, discrimination performance improves in some cases for biomarker combinations requiring the colocalization of two unique cancer-associated biomarkers in a biomarker-dependent manner (Figure 4D; middle panels). For example, MUC1 capture/FOLR1 + FOLR1 capture resulted in no discrimination of unspiked versus spiked plasma (ΔCt = 0.6) whereas MUC1 capture/BST2 + BST detection resulted in improved discrimination (ΔCt = 3.2). An assay configuration with all three biomarkers, MUC1 capture/FOLR1 + BST2 detection resulted in a ∼260,000 (2^18^) fold reduction of signal in the unspiked plasma sample, enabling the greatest separation of unspiked versus spiked plasma (ΔCt = 6.1, right panel).

### Biomarkers are colocalized on individual EVs from human cell lines

To confirm biomarker colocalization on individual EVs using an orthogonal approach, we utilized super-resolution microscopy to identify triple positive (BST2+/FOLR+/MUC1+) EVs. Figure 5 demonstrates the colocalization and simultaneous detection of FOLR1, BST2, and MUC1 on individual, representative COV413A cell line EVs captured on the surface of a microfluidic chip using phosphatidylserine directed S4 affinity from Oxford Nanotechnology (ONI, Oxford, UK).

**Figure 5:**
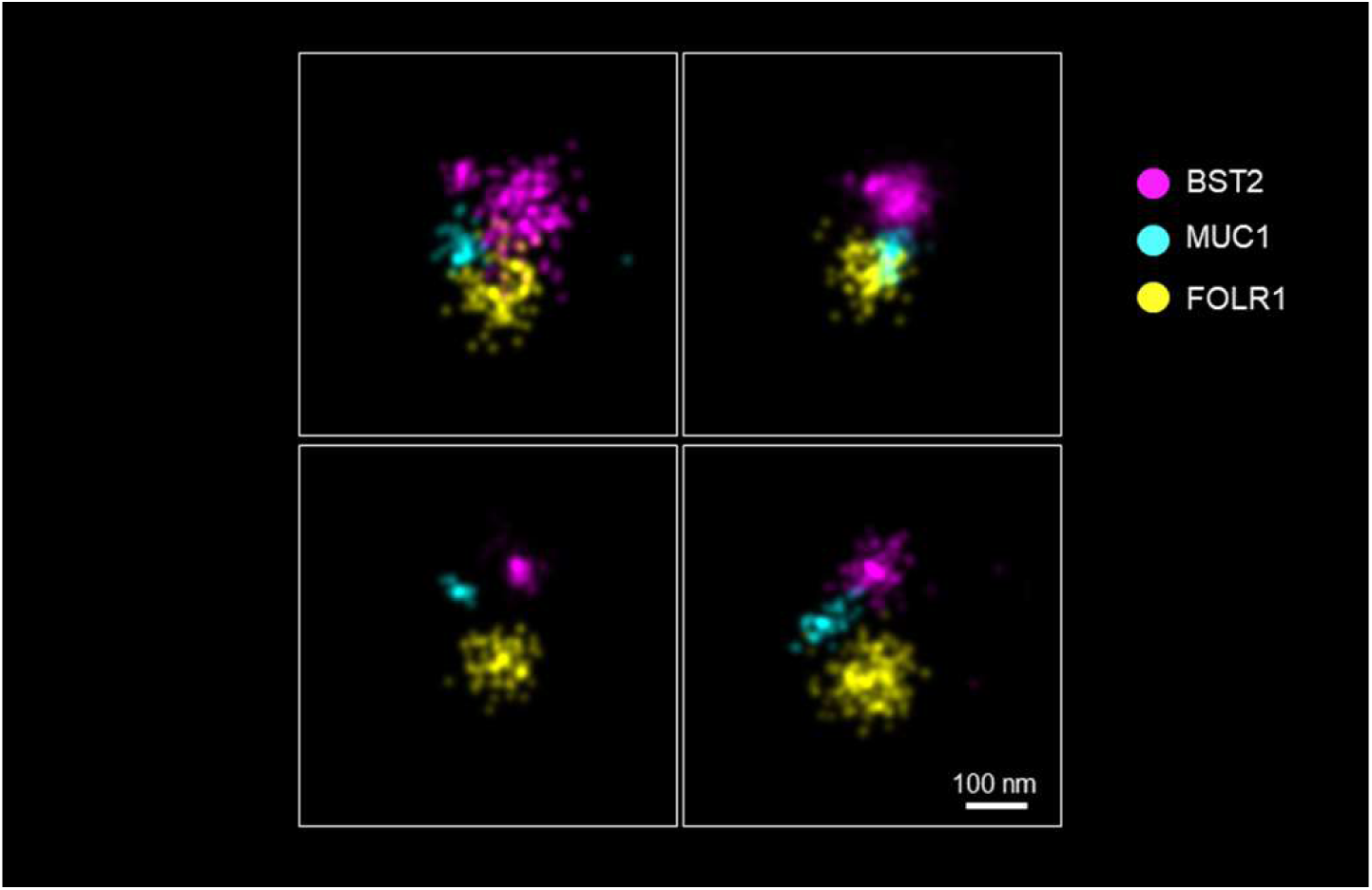
Colocalization of BST2, FOLR1, and MUC1 on single EVs using super-resolution microscopy. EVs derived from COV413A cells were captured on an S4 microfluidic slide (ONI) prior to staining with an antibody cocktail containing Alexa Fluor antibody conjugates to BST2 (magenta), MUC1 (blue), and FOLR1 (yellow). Triple-positive EVs were identified using CODI software and verified manually. Representative images are shown. The scale bar represents 100 nm.

### Assay performance depends on intact EVs

To demonstrate that assay signal is derived from colocalized biomarkers on an intact EV, SEC enriched EVs from human cell line conditioned media (COV413A) or pooled patient plasma from late-stage HGSOC were subjected to capture with MUC1 antibodies. Prior to detection, captured EVs were incubated with buffer alone (-) or buffer containing 0.075% Triton-×100 (+) sufficient to disrupt intact EVs (Supplemental Figure 4)(Osteikoetxea et al., 2015). Assay signal was generated using detection antibodies to BST2 and FOLR1. Detergent treatment results in a significant increase in Ct suggesting that most of the signal is derived from intact EVs that are detergent sensitive.

### Assay signal requires biomarker colocalization on the same EV

Theoretically, complementary DNA-conjugated antibodies located on adjacent EVs could generate a DNA template if within sufficient proximity (“bridging”). We characterized a mixture of EVs from two different cancer cell lines, OVKATE and SW900, each predicted to express high levels of two of the three biomarkers (BST2^high^/FOLR1^high^/MUC1^low^, and BST2^high^/FOLR1^low^/MUC1^high^, respectively). To assess bridging, we characterized the individual cell line EVs (4×10^8^ particles/mL) and a 1:1 mixture (2×10^8^ particles/mL each) using the triplet biomarker combination BST2 capture/FOLR1 + MUC1 detection). As expected, EVs from COV413A that express all three biomarkers produced high signal (18.4 Ct) relative to EVs from G-401 cells which express low levels of both FOLR1 and MUC1 (33.4 Ct). EVs from double-biomarker positive cell lines OVKATE (BST2^high^/FOLR1^high^/MUC1^low^) or SW900 (BST2^high^/FOLR1^low^/MUC1^high^) generated assay signals of 26.7 Ct and 30.5 Ct, respectively, while the 1:1 mixture of EVs from both cell lines generated a signal of 28.2 Ct (Supplementary Figure 5). The 1:1 mixture signal was consistent with the arithmetic mean of the individual cell line EVs, suggesting that bridging does not contribute significantly to assay signal.

### Assay linearity and reproducibility

The linearity of the MUC1, BST2+FOLR1 combination was evaluated using plasma spiked with a 4-fold dilution series of EVs collected from the human ovarian cancer cell line COV413A (see Supplemental Figure 6). Samples prepared in this manner were expected to cover the healthy control and ovarian cancer cycle threshold range of ∼20 to ∼40 Ct. The dilution series was prepared 4 times and then each sample was run in duplicate (8 replicates) in a single assay plate. There was no high-dose hook effect observed, even at the highest spike-in level of 3.2×10^8^ EVs per sample well, which is well above the combination Ct values observed in either healthy or ovarian cancer samples. This combination was linear over the 4-log range of 19,500 to 320 million cell line EVs per well. Based on the results from the linearity study 5-level controls were targeted at 3.2×10^8^ EVs/well, 8×10^7^ EVs/well, 2×10^7^EVs/well, 5×10^6^ EVs/well, and unspiked plasma background. Each of these full process controls was tested with the MUC1, BST2+FOLR1 combination on 16 separate days and the between-day reproducibility was calculated from these data. This combination had between-day coefficients of variation (CVs) of under 2% for all 5-level controls.

### Assay signal correlates with cell line gene expression and protein expression

To evaluate the correlation between assay signal and cell line gene and protein expression, purified EVs from 32 human cancer cell lines were characterized at a constant nanoparticle concentration using single-biomarker capture/detect combinations directed towards BST2, FOLR1, and MUC1. Assay signal for each biomarker is summarized in Supplemental Figure 7 along with published gene expression and protein expression data (when available). Signal strength was positively correlated with gene and protein expression for all three biomarkers (Spearman’s correlation coefficients between 0.47 and 0.87), suggesting that cell line biomarker expression is predictive of EV protein composition.

### Discrimination of high grade serous ovarian cancer (HGSOC) in patient samples

To assess feasibility for the detection of early-stage cancer, we applied the assay to a clinical cohort of 92 patients comprising 42 HGSOC cases (18 stage I/II, 24 stage III/IV), 26 females with benign adnexal masses, and 24 female healthy controls. Plasma from each subject was characterized using the BST2/FOLR1/MUC1 biomarker combination targeting BST2/FOLR1/MUC1-positive EVs. The results are summarized in Figure 6. In this limited proof of concept study, the biomarker combination of BST2/FOLR1/MUC1 achieved a 55.6% sensitivity (95% CI: 30.8%-78.5%) for stage I/II disease. More importantly, this biomarker combination readily discriminates between early-stage and benign disease. This performance has been further improved in subsequently developed assay configurations (data not shown), suggesting this approach may have clinical utility in detecting early stage HGSOC and discriminating ovarian malignancy from benign adnexal masses.

**Figure 6.**
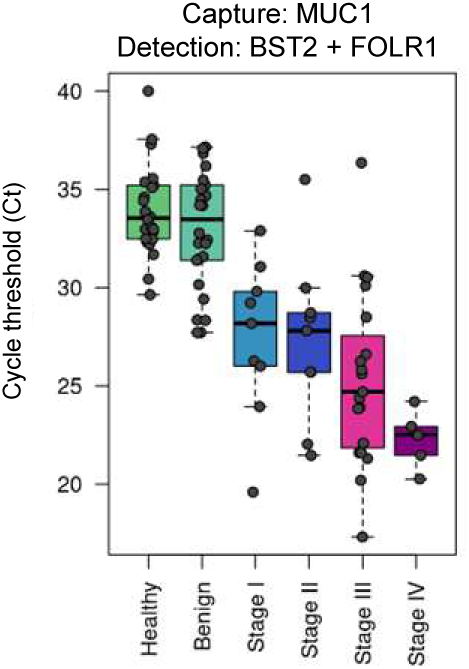
Discrimination of High-Grade Serous Ovarian Cancer (HGSOC) from Healthy Controls and Benign Ovarian Masses. Performance of the MUC1 capture/BST2 + FOLR1 detection biomarker combination in a cohort of healthy controls (n=24), patients with benign ovarian masses (n=26), and patients with stage I (n=9), II (n=9), III (n=15), and IV (n=4) HGSOC in a proof-of-concept assay.

## Discussion

Cancer screening has been shown to reduce cancer-associated mortality through the detection of localized disease that is more amenable to treatment, underscoring the clinical need for more accurate and accessible screening tests (Crosby et al., 2022). Despite considerable effort developing blood-based screening approaches, technical challenges have constrained test performance. Early-stage tumors are small, limiting the abundance of the available analytes for measurement in liquid biopsies. Cancer mis-regulates normal physiological processes, which means the analytes produced by cancers are often also produced by normal tissues, either in the tissue of origin or other parts of the body. Even when arising from the same tissue type, cancer utilizes blind evolutionary opportunism and therefore the mechanisms that lead to carcinogenesis can vary widely. Measuring colocalized cancer-associated biomarkers on intact EVs addresses each of these challenges. Specifically, EVs are a stochastic sampling of the cell of origin bearing cancer-associated biomarkers both on their surface and as cargo (Doyle and Wang, 2019; Li et al., 2020). They are released continuously from tumor cells due to their anti-inflammatory and tumor-supporting functions and are stable enough to tolerate processing and amenable to at-home collection tests (Johnsen et al., 2019).

Here we describe a novel, non-invasive method for the early detection of cancer through the measurement of co-localized cancer associated biomarkers on the surface of extracellular vesicles. EVs are an emerging analyte class with features well suited to use in diagnostic testing and represent an area of science that is undergoing rapid advances in the understanding of basic biology as well as research and clinical applications (Ghodasara et al., 2023). There are biological and technical limitations to the approach described here that are important considerations informing ongoing assay development. EVs derived from hematopoietic lineage cells comprise 99.8% of all EVs in human plasma, and thus biomarker targets that are highly expressed in blood may generate assay background (Auber and Svenningsen, 2022; Ferguson and Weissleder, 2020; Johnsen et al., 2019). Assay performance is dependent on the quality and availability of compatible affinity reagents, such as antibodies. Promising cancer-associated biomarkers lacking affinity reagents with suitable performance characteristics may restrict potentially useful biomarker combinations from inclusion in assays. Among biomarkers with suitable affinity reagents, we observed a range in discrimination performance, potentially resulting from variability in healthy background signal. Variability in assay specificity may be driven by several factors including biomarker expression, antibody affinity and specificity, and the orientation of the assay design (i.e., the choice of biomarkers for use in immunoaffinity capture versus the detection step). The studies we conducted evaluating test performance in clinical samples provide encouraging evidence that the assay will have utility in identifying early-stage cancers and discriminating malignant from healthy samples but represent proof of concept only. Further studies to assess the clinical performance of the assay for the detection of high-grade serous ovarian cancer (HGSOC) in well powered case-control studies utilizing clinical samples collected prospectively under standardized protocols, are in progress.

Measuring colocalized biomarkers on single EVs using the assay described herein is a promising approach for early-stage cancer detection. It addresses the biological abundance limitations of genomic liquid biopsy approaches while offering improved specificity over bulk EV measurements. The assay measures EV surface markers and thereby leverages the natural enrichment of membrane-bound epitopes relative to cargo proteins due to the 100-fold increase in surface area-to-volume ratio between an EV and its cell-of-origin. Moreover, it can interrogate billions of EVs simultaneously enabling the detection of dilute (<1 in 100,000)(Ferguson and Weissleder, 2020) cancer associated EVs reaffirming the observation that the assay can be used to detect EVs from early-stage tumors. The qPCR-based readout enhances assay technical sensitivity relative to conventional enzyme-amplified sandwich immunoassays, similar to other approaches using immuno-PCR (Niemeyer et al., 2005; Stiller et al., 2021), proximity ligation assays (Tavoosidana et al., 2011; Weibrecht et al., 2010), and proximity extension assays (Larssen et al., 2017; Lundberg et al., 2011).

The small sample volume requirement, compatibility with standard blood collection tubes, and use of low-cost qPCR for signal generation are features that may help to enable broad adoption, including in lower resource settings. Advances in affinity reagent development and EV biomarker discovery may be leveraged to improve existing assays by incorporating new reagents or designing orthogonal combinations to boost assay performance. We anticipate this assay format is readily extensible to a broad array of clinical indications, across cancer types and beyond cancer. For example, neuronally derived EV biomarkers have been reported to track with cognitive decline in Alzheimer’s disease and may represent another opportunity for diagnostic utility (Eren et al., 2022). EV-based measurement of tumor-derived biomarkers may be helpful in guiding clinical management through use in companion diagnostic applications. For example, folate receptor alpha, one of the ovarian cancer biomarkers included in the combination described in this study, is the target of several ongoing efforts focused on the development of antibody-drug-conjugate (ADC) therapeutics (El Bairi et al., 2021). A liquid biopsy assay indicative of the presence of folate receptor alpha positive tumors may be helpful in identifying patients more likely to respond to targeted ADC therapy. The EV-based test method described here has the potential for immediate clinical utility in single cancer early detection, and potentially for multi-cancer early detection either through the rational combination of single-cancer EV tests, or as part of an integrated multi-omic approach.

## Materials and Methods

### Cell culture

All human cancer cell lines used in this study, and their respective sources, are indicated in Supplemental Table 1. SK-MEL-1, and SK-MES-1 were used under license from the Memorial Sloan Kettering Cancer Center. NCI-H146, NCI-H441, NCI-H520, and NCI-H1781 were developed by Dr. Gazdar and Dr. Minna and used under license from the National Cancer Institute. COV362 and COV413A were used under license from the European Collection of Authenticated Cell Cultures (ECACC). OVSAHO and OVISE were obtained from Sekisui XenoTech (Kansas City, KS, USA) under license from the Japanese Collection of Research Bioresources (JCRB, Osaka, Japan). All cells were thawed, subcultured, and maintained in complete growth media as recommended by the supplier. For EV production, cells were passaged into large capacity tissue culture-treated flasks or dishes until 70-80% confluent. The flasks or dishes were then transitioned to EV-depleted FBS (A2720801, ThermoFisher Scientific, Waltham, MA) or serum-free Cancer Cell Line Media XF (CCLM XF, C-28077, PromoCell GmbH, Heidelberg, Germany), depending on the individual cell line as follows; growth media was removed and the cells were rinsed twice with 1/2 volume of either complete media supplemented with exosome-depleted (EV-) FBS or CCLM XF serum free media prior to being replaced in basal media with 10% EV-FBS, or CCLM XF. After 48 hours, the conditioned media (CM) was collected and cellular debris removed by centrifugation at 300×*g* for 5-minutes followed by two subsequent centrifugations at 1500×*g* for 10 minutes each. All centrifugations were done at 21°C. The cleared CM was transferred to 50 mL conical tubes in ∼45 mL aliquots and stored at -80°C prior to purification.

### Concentration of conditioned media

Typically, 200 mL of CM was thawed at room-temperature and centrifuged at 1,300×*g* for 10 minutes to remove any insoluble debris. The supernatant was transferred via serological pipet to fresh 50 mL conical tubes then re-centrifuged at 2,000×*g* for 30 minutes. The supernatant was carefully transferred to a fresh tube and concentrated using Amicon Ultra Centrifugal filter units with a 10,000 molecular weight cut-off (Millipore-Sigma, St. Louis, MO). This process was repeated until all the retained material was concentrated to approximately 1 mL. All centrifugations were done at 21°C unless specified otherwise.

### EV Isolation from concentrated conditioned cell culture media or plasma

EVs were purified from either 1 mL of concentrated CM, acid-citrate-dextrose (ACD) plasma, or human potassium EDTA plasma. In each instance, 1 mL of concentrated CM or plasma was applied to an Izon qEV original 70 nm Legacy SEC column (ICO-70, Izon Science US, Medford, MA, USA) following the vendor-recommended protocol. Briefly, qEV columns were allowed to equilibrate to room temperature and washed in 10 mL room-temperature 1x PBS. Concentrated CM (described above) was applied to the top of the column and the flow through material was discarded. Two milliliters of 1X PBS were applied to the column and collected as the pre-EV fraction and analyzed as described. EV-elution fractions (EV1, EV2…EV8) were then collected by applying 0.5 mL 1X PBS sequentially until all desired fractions were collected. The PCV was determined based on physical and functional characterization of each EV fraction. Purified plasma EVs were immediately analyzed using the assay as described. Concentrated CM EVs were quantified using ZetaView Classic S model nanoparticle tracking analysis (NTA) instrument (Particle Metrix, Amersee, Germany) prior to aliquoting and freezing at -80^○^C. The protocol for purifying plasma and CM EVs was identical, with the exception that 0.02 uM-filtered 1X PBS, pH 7.4 was used for the concentrated CM EV purification (so particles in the PBS will not confound the downstream quantification) while unfiltered 1X PBS, pH 7.4 was used to purify EVs from plasma.

### Western blot analysis of human cell line or plasma extracellular vesicle fractions

Whole-cell extracts (WCE) of COV413A cells were prepared by scrape-harvesting 90%-95% confluent cells in 1X #9803 Cell lysis buffer (Cell Signaling Technology, Danvers, MA). Lysates were incubated at 4^○^C for 30 minutes prior to ultrasonic cavitation (3 x 15 seconds at 33% power) using QSonica Q125 Ultrasonic Disruptor (QSonica, Newtown, CT, USA) fitted with a 2 mm titanium probe tip. Protein concentrations were determined, and the lysate was further diluted to 1mg/mL in 4X LDS Sample Buffer (NP0008, ThermoFisher Scientific). Human plasma lysate was prepared by mixing whole human plasma 1:1 with 4X LDS Sample Buffer followed by ultrasonic cavitation to homogenize the sample. Individual EV fractions from concentrated CM were diluted 1:1 in 4X LDS Sample Buffer and human plasma EVs were concentrated 10:1 prior to 1:1 dilution in 4X LDS Sample Buffer. In all cases, samples were heated to 70^○^C for five minutes prior to use. For polyacrylamide gel electrophoresis (PAGE) and western blot experiments, 10-20 µL of WCE, plasma lysate, or EV fraction was loaded alongside a biotinylated ladder and/or pre-stained protein molecular weight markers (Catalog numbers 7727 and 59329, respectively, Cell Signaling Technology) on NuPAGE™ Bis-Tris™ pre-cast gels (ThermoFisher Scientific). Gel percentage, running buffer, and transfer conditions were dictated by the apparent molecular weight of the proteins being immunoblotted. In most cases, we used 1.0 mm, 4-12% Bis-Tris mini gels with MOPS-based running buffer (NP0001, ThermoFisher) according to the manufacturer’s instructions. After PAGE, gels were either stained with Gel-Code™ Protein Stain (24590, ThermoFisher) or transferred to 0.2-micron PVDF membranes (1620174, Bio-Rad, Hercules, CA, USA) using a XCell SureLock Mini-Cell (EI9051, ThermoFisher) according to the manufacturer’s recommendations. After electrophoretic transfer, all membranes were blocked at room temperature for one hour in 1X TBST (#9997, Cell Signaling Technology) supplemented with 5% non-fat dry milk (AB10109, American Bioanalytical, Canton MA). Primary antibodies (Supplemental Table 1) were incubated overnight (16 hours) in 1X TBST supplemented with either 5% non-fat dry milk or bovine serum albumin (#9997, #9998, Cell Signaling Technology) at 4^○^C with constant rocking according to the manufacturer’s recommendation. Each blot was washed thrice for 5 minutes in 1X TBST at room temperature prior to addition of HRP-conjugated secondary antibodies and HRP-anti-biotin antibody, both diluted in 1X TBST + 5% non-fat dry milk, for one hour at room temperature. Each blot was then washed thrice in 1X TBST for 5 minutes and proteins were detected using either LumiGLO/Peroxide Reagent or SignalFire™ ECL reagent (7003 and 6883, respectively, Cell Signaling Technology) and an iBright™ imaging system (ThermoFisher). All images are presented with no image adjustments (other than cropping to highlight the area of interest) and are typically between 5 and 15-minute exposures, depending on the signal strength. All antibodies used for western blot and their working dilution are listed in Supplemental Table 1.

### Characterization of EV size distribution and concentration

Size distribution and particle counts were determined using a ZetaView NTA instrument after enrichment of the EVs by SEC. ZetaView software 8.05.14 SP7 (Particle Metrix) acquired and exported the data. The instrument was calibrated as recommended by the manufacturer prior to all measurements. Following SEC isolation and prior to NTA measurement, the isolated particles were diluted 1:250 - 1:2000, which allowed the instrument to accurately measure the particle distribution (50-200 particles per field of view). As recommended by the manufacturer for the quantification of EVs, the sensitivity was set at 80, shutter at 100, min brightness 20, min area 10, and max area 10000. All 11 positions of the instrument were analyzed, with outlier positions excluded automatically by the instrument. The size distribution results were binned in 1nm intervals from 1 to 1000 nm.

### Antibody Panels

All antibodies used for assay development were obtained from commercial sources (Supplemental Table 3). Initially, around 50 different antibodies against the three biomarker targets were screened for functionality and compatibility using cell line EVs. Antibody clones that passed primary screening were further optimized to determine the optimal conjugation concentration of both probes and beads in biomarker-expressing cell lines vs biomarker-negative cell lines or buffer alone controls. The top antibodies for the MUC1, BST2, and FOLR1 combination were then validated in human plasma and further down-selected based on those antibodies that produced the best discrimination of HGSOC from healthy women and those with benign adnexal masses.

### Plasma Samples

The 124 healthy donor plasma samples were collected under a WCG IRB-approved protocol (20212722). Each donor provided 2 tubes of blood which were collected using a 10-mL collection volume K_2_EDTA tube (0265732, Thermo Fisher Scientific). After collection, each tube of K_2_EDTA plasma was inverted 6-8 times, centrifuged (1500×g at room temperature for 15 minutes) at the collection site within 60 minutes collection to separate the plasma from the other blood components. The resultant plasma was divided into 1 mL aliquots in 2 mL cryovials, frozen and stored at -80°C. The frozen aliquots were shipped from the collection site to Mercy BioAnalytics on dry ice and stored at -80°C. A plasma pool from ninety healthy women was prepared in-house using samples purchased from ProteoGenex Inc. (Inglewood, CA). Late- and early-stage HGSOC cases were purchased from ProteoGenex, Inc. and Reprocell (Beltsville, MD). All ProteoGenex plasma samples were sourced from Moscow, Russia and collected in either K_2_EDTA or K_3_EDTA blood collection tubes. Reprocell, Inc. samples were sourced from the United States and collected in acid citrate dextrose (ACD) blood collection tubes. Plasma samples from women with benign ovarian masses were purchased from ProteoGenex were also sourced from Moscow, Russia and collected in K_2_EDTA blood collection tubes. Patient sample attributes are shown in Supplemental Table 4.

### Antibody-functionalized magnetic beads

Antibodies containing bovine serum albumin (BSA) were purified using protein A and G Spintrap purification kits (GE28-9031-34 or GE28-9031-32, Cytiva, Marlborough, MA) according to manufacturers’ instructions. All antibodies used in the assay underwent a buffer exchange process to ensure that any carriers or impurities that might interfere with conjugation were removed. Up to 500 µg of antibody material was added to 2 mL of sterile 1X PBS (10010031, ThermoFisher Scientific) in 30K MW Amicon Ultra Filter Units (UFC203024, Sigma-Aldrich, St. Louis, MO, USA) and centrifuged for ten minutes at 4,375×*g*. This process was repeated 6 times, after which the antibodies were recovered and sterile filtered by centrifugation in 0.22 µM Spin-X tube filters (8160, Corning-Costar, Corning, NY) for 2 minutes at 12,000×*g*. Antibody concentrations were measured using the NanoDrop spectrophotometer. Immunoaffinity magnetic beads were prepared by coating antibodies onto Dynabeads® M-270 Epoxy beads using the Dynabeads Antibody Coupling Kit (14311D, ThermoFisher Scientific) according to the manufacturer’s instructions. Briefly, the beads were suspended at 10 mg mL^-1^ in C1 buffer and mixed thoroughly by vortexing. 1 mL aliquots of the beads were transferred to 1.5 mL tubes, which were then placed on a DynaMag™-2 magnetic rack (12321D, ThermoFisher) for 1 minute to separate the beads from the solution and the supernatant was removed. C1 buffer and 20-160 µg of capture antibody, totaling a volume of 500 µL, were added and the beads were resuspended by gentle vortexing. After adding 500 µL C2 buffer, the beads were gently vortexed, and the tubes were placed on a HulaMixer™ (ThermoFisher Scientific) for conjugation at 37°C for 19 ± 0.5 h. The tubes were briefly centrifuged at 300×*g* for 10 seconds, placed on the magnetic rack for 1 minute to separate the beads from the solution, and the supernatant was removed. The beads were washed using four cycles of resuspension and pulse centrifugation in Dynabeads wash solutions containing 0.05% Tween 20 (H5152, Promega, Madison, WI, USA), followed by three washes in 1X PBS (10010072, ThermoFisher Scientific) with 0.1% BSA (0332, VWR, Radnor, PA, USA) and 0.1% Pluronic F-68 (P1169, Spectrum Chemical, New Brunswick, NJ, USA). The antibody-functionalized magnetic beads were resuspended in 1 mL of 1X PBS, pH 7.4 with 50% glycerol (49767, Sigma-Aldrich), 0.1% BSA, and 0.1% Pluronic F-68 and stored at -20°C.

### DNA-conjugated antibody preparation

Antibodies were buffer exchanged and purified, if needed, as described above. Antibodies were subsequently conjugated to DNA using a copper-free click chemistry reaction. Oligonucleotides (Supplementary Table 2) were synthesized and purchased as single-stranded DNA (ssDNA) from Integrated DNA Technologies, Inc. (Coralville, IA, USA) or Bio-Synthesis, Inc. (Lewisville, TX, USA). Before performing the conjugation, the ssDNA was annealed to form a double-stranded product. Complementary ssDNA strands (Strands 1+3 and Strand 2+4, Table 1) were mixed to a final concentration of 50 µM in 1X PBS and annealed using a standard thermal cycler (Applied Biosystems, ProFlex PCR System). The annealing protocol was as follows: 95°C for 2 minutes, 90°C for 3 minutes, and then reducing the temperature by 5°C in 3-minutes intervals until 25°C was reached. Monoclonal antibodies at 1 mg mL^-1^ in 1X PBS were incubated with DBCO-PEG5-NHS Ester linker (A102P, Click Chemistry Tools, Scottsdale, AZ, USA) at a 4:1 molar ratio of crosslinker to antibody for at least 2 h at room temperature. Following incubation, the antibody-linker solution was washed once using a 30 kDa Amicon Ultra-2 centrifugal filter (UFC203024, Sigma-Aldrich). Annealed DNA oligonucleotides with a 5′ azide modification (Integrated DNA Technologies) were added to the activated antibodies at a 3:1 molar ratio of oligonucleotide to antibody and were allowed to react overnight at room temperature. The conjugate concentrations were measured using the Qubit® protein assay kit (Q33211, ThermoFisher Scientific) according to the manufacturer’s instructions, and the DNA-conjugated antibodies were stored at 4°C in 1X PBS.

**Table 1:**
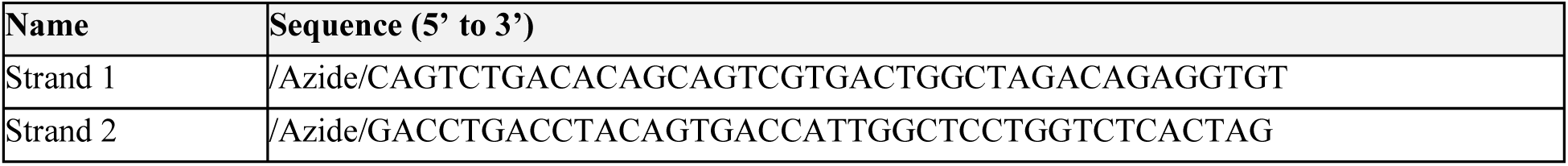
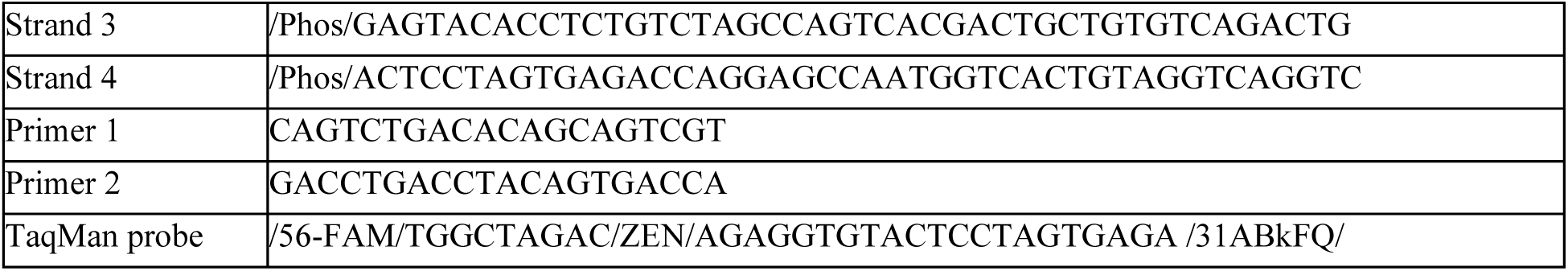
Oligonucleotide Sequences for Primers and TaqMan Probes.

### Assay protocol

Following EV isolation by SEC, a concentrated buffer solution was added to the purified EVs to obtain working concentrations of 0.1% BSA, 0.1% Pluronic F-68, 100 µg mL^-1^ salmon sperm DNA (15632011, ThermoFisher), and 0.1% ProClin 950 (46878-U, Sigma-Aldrich). Antibody-functionalized magnetic beads directed against Biomarker 1 were diluted 1:2 with 0.1% BSA, 0.1% Pluronic F-68, 100 µg mL^-1^ salmon sperm DNA, and 0.1% ProClin 950, and 4.8 µL aliquots of the diluted beads per well (corresponding to 16 µg of beads/well) were added to the wells of a 96 deep-well KingFisher plate. 250 µL of the diluted, purified EVs was added to each well of a deep-well KingFisher plate. The sample plate was covered with an adhesive plastic film, was placed on an Eppendorf ThermoMixer C (Eppendorf, Enfield CT, USA) and incubated at 21°C for 2 hours, mixing at 1100×g During this incubation, all the wash and reaction plates required for the assay were prepared and are described in Table 2:

**Table 2:** Configuration and Composition of the 96-Well KingFisher Plates.

| Reaction plate type | # | Volume/well | Components |
| --- | --- | --- | --- |
| Antibody plate | 1 | 100 $\mu\text{L}$ | DNA-conjugated antibody, 0.1% BSA, 0.1% Pluronic F-68, 100 $\mu\text{g mL}^{-1}$ salmon sperm DNA |
| TEPBS wash | 2 | 1 mL | 1X PBS, pH 7.4, 0.05% Tween 20, 5 mM EDTA |
| PBST wash | 1 | 1 mL | 1X PBS, pH 7.4, 0.05% Tween 20 |
| Ligation plate | 1 | 100 $\mu\text{L}$ | 75,000 units $\text{mL}^{-1}$ of T7 DNA ligase (New England Biolabs, M0318), 33 mM Tris-HCl, 5 mM $\text{MgCl}_2$, 0.5 mM ATP, 0.5 mM DTT 3.75% PEG 6000, 0.5% Tween 20 |
| PBST transfer plate | 1 | 150 $\mu\text{L}$ | 1X PBS, pH 7.4, 0.05% Tween 20 |
| qPCR plate | 1 | 30 $\mu\text{L}$ | TaqMan Fast Advanced Master Mix (4444557, Applied Biosystems), 500 nM primers, 200 nM TaqMan probe |

After the 2-hour EV-bead-binding incubation was complete, a KingFisher Flex System (ThermoFisher Scientific) was loaded with the sample plate, the antibody-probe plate, the wash plates, the ligation reaction plate, and the PBST transfer plate and the custom KingFisher program initiated. Briefly, the beads are magnetically captured and incubated in the antibody-probe plate for 32 minutes and 50 seconds, then washed twice in TEPBS for three minutes and 30 seconds, then washed in PBST for three minutes and 30 seconds and placed in the ligase reaction plate for 21 minutes and 40 seconds. Finally, the beads are transferred into a shallow plate containing PBST. The beads are then manually transferred to a qPCR plate using a KingFisher Flex 96 PCR magnetic head. qPCR plates were covered with a clear plastic adhesive and placed into QuantStudio Dx Real-Time PCR instruments (ThermoFisher Scientific) run using the following amplification conditions: 50°C for 2 minutes, 95°C for 2 minutes, 40 cycles at 95°C for 5 seconds, and 60°C for 20 seconds. See Table 2 for primer and TaqMan probe sequences. All samples were run in either duplicate or triplicate and those showing a Ct spread >3 among the replicates were excluded from subsequent data analysis.

### Detergent disruption of EVs

To demonstrate the assay is EV-based, a detergent-sensitivity experiment was performed. Exposure of EVs to detergent would disrupt their membranes, resulting in loss of MH assay signal. As described in the paragraph above, EVs purified with SEC were diluted in a buffer containing 0.1% BSA, 0.1% Pluronic F-68, 100 µg mL^-1^ salmon sperm DNA, and 0.1% ProClin 950. Antibody-functionalized beads were used to capture the purified EVs in a 2-hour incubation under agitation. The beads with the captured EVs were then transferred, using the KingFisher Flex system, to deep-well plates containing 0.1% BSA, 0.1% Pluronic F-68, 100 µg mL^-1^ salmon sperm DNA, 0.1% ProClin 950, and 0.075% Triton X-100 (T8787, Sigma-Aldrich). The plates were incubated for 30 minutes, at 21°C, under agitation (1100×g) in a ThermoMixer C instrument to allow for EV lysis. After incubation with detergent, the deep-well plate containing the beads were washed and read out with detection antibodies to either tetraspanins or the indicated biomarkers as described above (Assay Protocol).

### Transmission Electron Microscopy of cell-line and plasma EVs

Cancer cell-line and HGSOC cancer EVs were prepared as described above and sent to Alpha Nano Tech (400 Park Offices Drive, STE 108, Morrisville, NC, 27709) in 1X PBS, on dry ice, for transmission electron microscope imaging using a Gatan Orius SC1000 CCD camera with Gatan Microscopy Suite 3.0 software. Copper-carbon formvar grids were glow discharged immediately prior to loading with the sample. Samples were processed undiluted. Grids were floated on 10 µL sample drop for 10 minutes, washed two times with water by floating on the drop of water for 30 seconds, and negatively stained with 2% uranyl acetate by floating on the drop of stain for 30 seconds. The grid was blot dried with Whatman paper and imaged with Jeol 1230 electron microscope. Pixel sizes for 40,000X, 80,000X, and 120,000X are 1.28 nm, 0.7 nm, and 0.47 nm, respectively.

### Super-resolution microcopy

EV biomarker colocalization was evaluated with direct Stochastic Optical Reconstruction Microscopy (dSTORM) using an ONI Nanoimager S and an EV Profiler Kit (Oxford Nanoimaging Inc., Oxford, UK). Antibodies against BST2, FOLR1, and MUC1 were conjugated to fluorophores using the Zip Alexa Fluor™ Rapid Antibody Labeling Kit from ThermoFisher Scientific with Alexa Fluor 488 (Z11233), Alexa Fluor 647 (Z11235), or Alexa Fluor 555 (Z11234), respectively. EVs collected from conditioned media from COV413A cells were purified by SEC as described above, characterized by NTA, then captured on imaging slides using S4 protein phosphatidylserine-dependent, protein-independent capture following the provided protocol (EVProfiler Kit v 1.5). Briefly, the microfluidic imaging chip was functionalized with S4 to capture phosphatidylserine containing vesicles then incubated with 5.6×10^8^ COV413A-derived EVs. After allowing the EVs to bind the surface, an antibody cocktail consisting of ONI N1 solution and 1X PBS containing 0.05% Tween with MUC1-AF647 (1:200), FOLR1-AF555 (1:500), and BST2-AF488 (1:50), was used to stain S4-bound EVs. After washing, the chip was prepared and imaged. A total of 7500 frames at 30 ms exposure time were collected with 1500 frames at 640 nm, 3000 frames at 561 nm and 3000 frames at 488 nm at 30%, 50% and 50% laser power respectively. The CODI software package (https://alto.codi.bio/, release date June 2023) was used for subsequent image analysis. EVs that were triple positive for MUC1, BST2, and FOLR1 were defined as clusters identified by density-based clustering algorithm (DBSCAN) with a radius of ≤80 nm and with ≥8 localizations of each biomarker.

### Statistical Analysis

Data is presented as PCR cycle threshold (Ct). When assay data was collapsed to a single value, the median of replicates was displayed. Cell line gene expression and mass spectrometry data was obtained from the Cancer Cell Line Encyclopedia (CCLE). Mass spectrometry data was not available for all cell lines. Heatmap data are ordered by signal strength. Heatmaps containing multiple data types were row-scaled with a cap of +/- 3 standard deviations to enable comparisons between different data modalities. When only gene expression was shown, heatmaps were ordered to match the cell lines displayed in the figure, log2 transcripts per million (TPM) displayed, and no scaling was used. Correlation analysis between assay signal and gene expression or mass spectrometry data was done using Spearman’s Rank Correlation. Assay sensitivity at 95% specificity was calculated using the epiR R package (version 2.0.40) and ROC curves were generated with the R package ROCit (version 2.1.1). All analyses were performed in R v.4.0.5.

## Acknowledgements

The authors would like to thank Maciej Pacula (Flagstaff Solutions, Acton, MA), David Ransohoff (UNC School of Medicine, Chapel Hill, NC), Steven Skates (Mass General Hospital, Harvard University, Boston, MA), Floyd Taub (Golden, CO), Daria Filonov (AlphaNano Tech, Durham, NC), and Ed Ha (Iskuda Therapeutics, New York, NY) for their generous support during the design, execution, and analysis of this work.

## SUPPLEMENTAL FIGURES

**Supplemental Figure 1:**
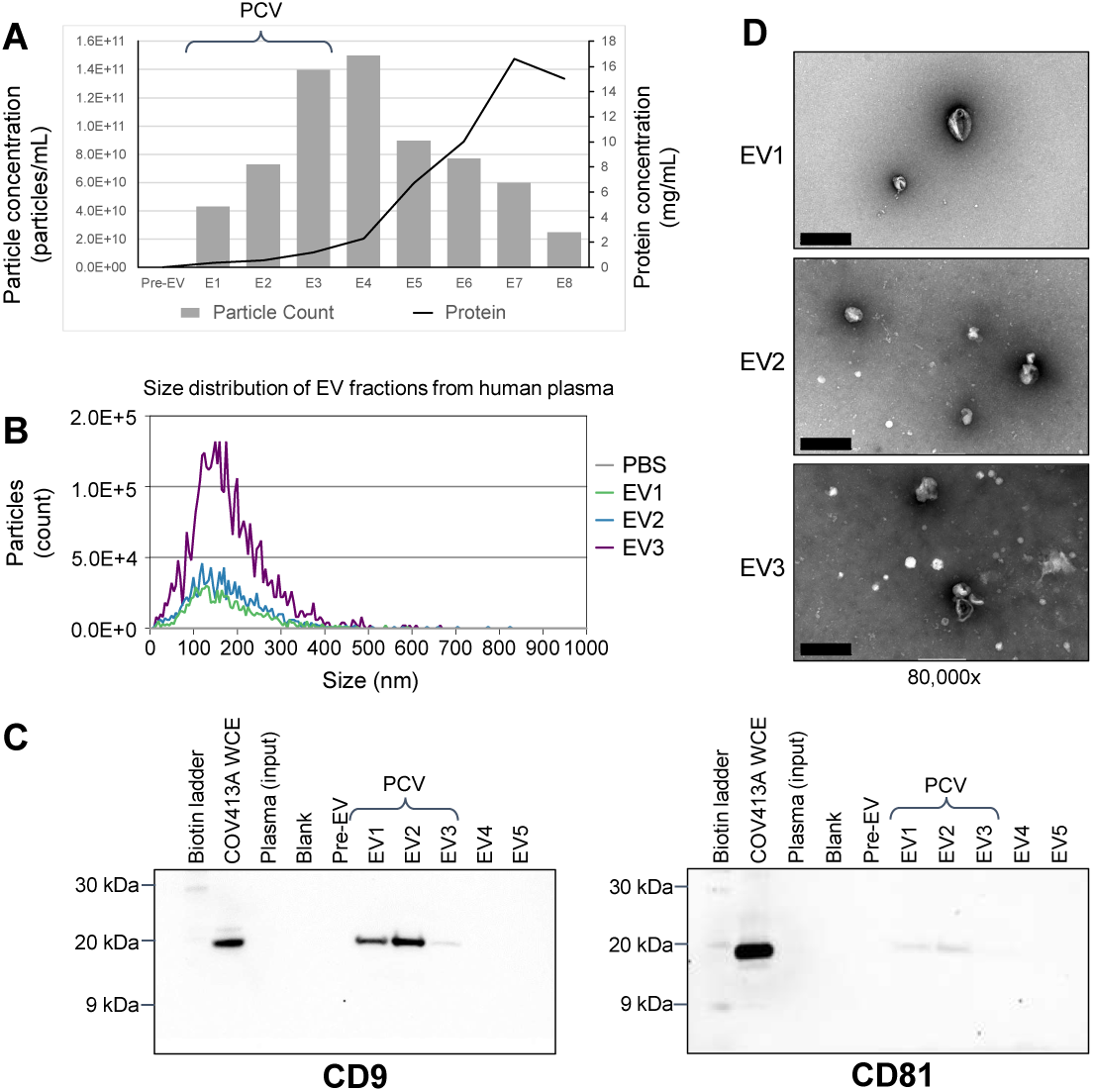
Characterization of human plasma EVs. EVs were enriched from a pool of healthy human plasma samples by SEC. Individual fractions were collected and subject to NTA. Unconcentrated SEC fractions (EV1, EV2, EV3) were characterized to determine particle counts (A) and overall size distribution (B) of the enriched nanoparticles. (A) Plot showing nanoparticle counts (grey bars) and protein concentration (black line) for the EV-containing SEC fractions from human plasma. The pooled collection volume (PCV), consisting of fractions EV1-EV3, used in the downstream assay, are highlighted. The Pre-EV sample is a sum of the void volume and the wash fraction containing lipoproteins and larger, non-EV particles (data not shown). (B) NTA size-distribution plot of individual fractions EV1, EV2, and EV3 demonstrating particles within the size range of 40-400 nm. Buffer alone (PBS) was used to determine the background. (C) In parallel, concentrated SEC fractions were analyzed by western blot against normal human plasma (input) and COV413A whole cell extract (WCE) as a positive control. CD9 (left panel) and CD81 (right panel) are undetectable in the input sample due to the low relative abundance in human plasma. The PCV is indicated demonstrating CD9 and CD81 presence in fractions EV1-EV3. (D) Representative 80,000x TEM images of EVs present in each of the three PCV fractions (EV1, EV2, and EV3) are shown, indicating intact particles between 40 and 400 nm. Scale bars are 500 nm.

**Supplemental Figure 2:**
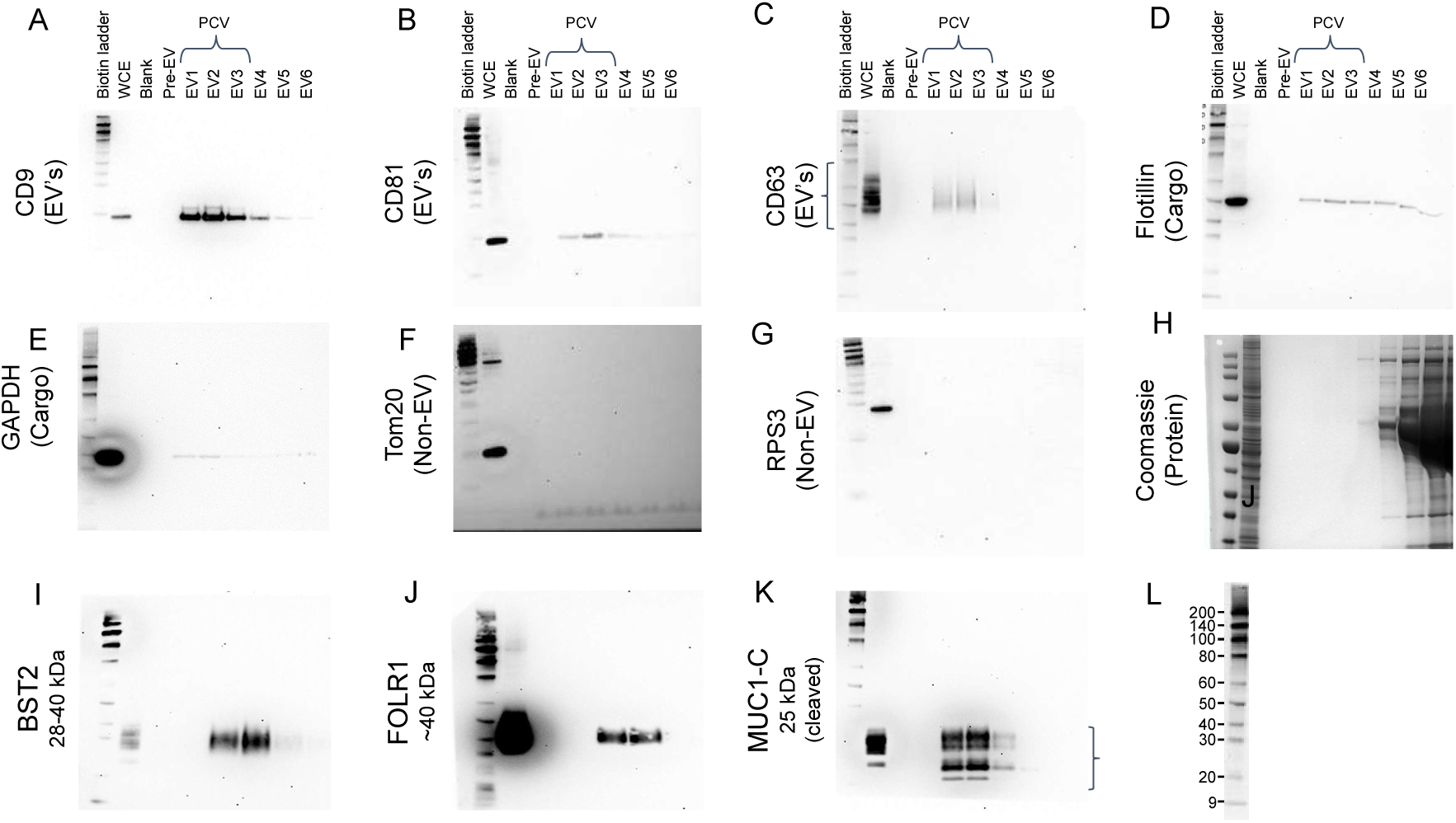
Expression of EV and Ovarian Cancer Biomarkers in EV Fractions. Full-panel blots of data shown in Figure 2 and additional western blot analysis of whole cell extract (WCE) from COV413A cells and SEC purified fractions from COV413A conditioned media (EV1-EV6) using antibodies against EV surface biomarkers CD9 (A), CD81 (B), and CD63 (C); EV cargo biomarkers Flotillin (D) and GAPDH (E) to demonstrate the EVs were intact EVs during enrichment; and non-EV internal membrane proteins Tom20 (F) and RPS3 (G) to demonstrate the EV fractions are the result of EV biogenesis and do not include intracellular vesicles. Total protein staining (Coomassie) was used to validate SEC separation of the EV fractions from the soluble protein component and was confirmed by measuring the protein concentration in each fraction (Figure 2). (I, J, K) Full panel blots of data shown in Figure 2 demonstrating abundance of BST2 (I), FOLR1 (J), and MUC1-C (K) in COV413A EVs. Representative molecular weight makers with indicated observed molecular weights are shown in L.

**Supplemental Figure 3:**
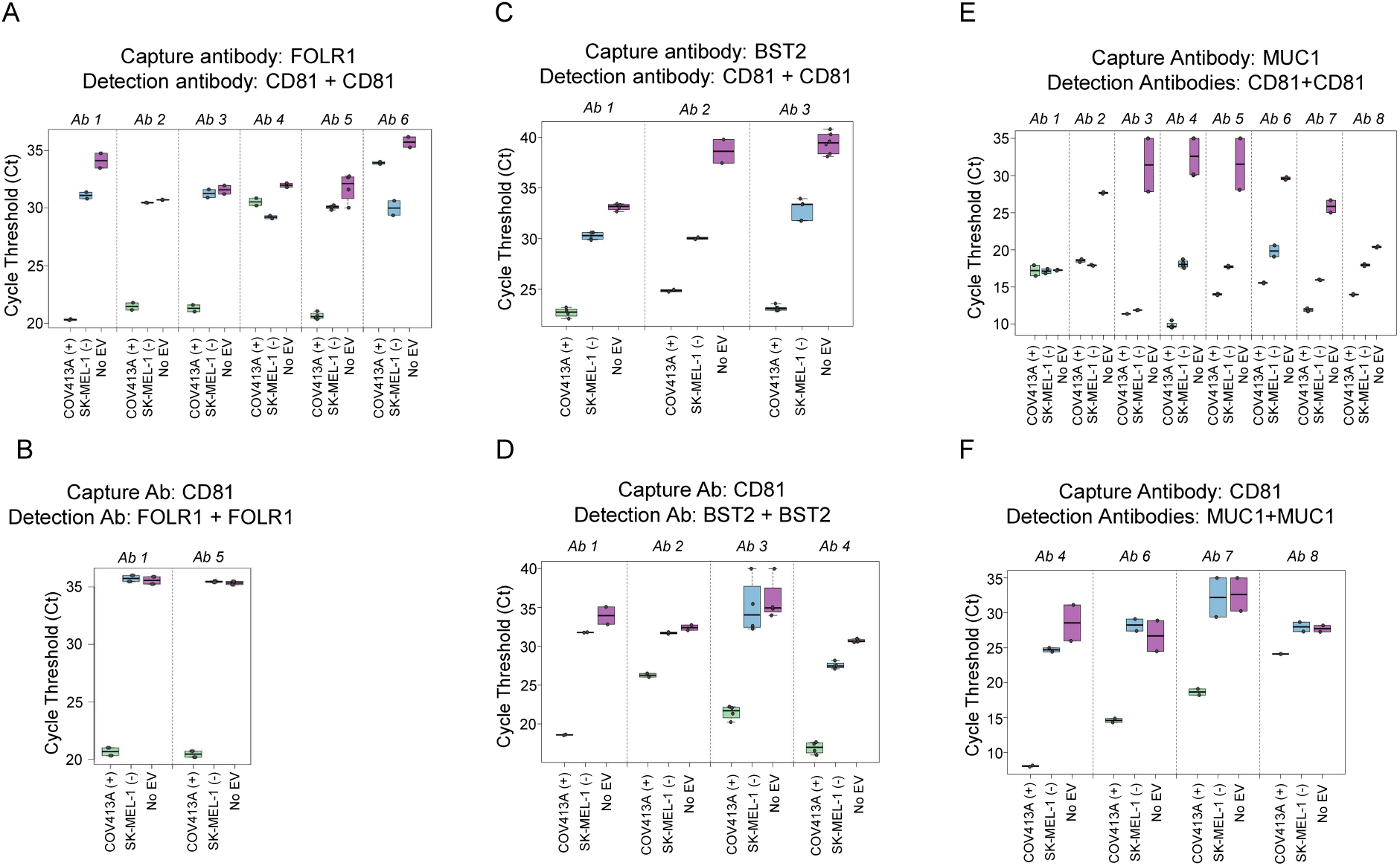
Screening antibodies for functionality and specificity for capture and detection of EVs. Distinct monoclonal antibody clones against FOLR1 (A, B), BST2 (C, D), and MUC1 (E, F) were screened for their ability to capture (A, C, E) and detect (B, D, F) SEC-purified EVs from human cell lines known to be positive (COV413A (+)) or negative (SKMEL1 (-)) for FOLR1, BST2, and MUC1. (A, C, E) Each antibody was independently conjugated to magnetic beads at a fixed density and used to capture cell line EVs in solution. Captured EVs were detected using complementary anti-CD81 dsDNA-conjugated detection antibodies. Antibody clones demonstrating both functionality (greatest signal to noise between the positive and negative cell line) and specificity (low Ct in positive cell line and high Ct in negative cell line) were further tested and optimized (not shown). No EV controls were used to eliminate cross-reactive or “sticky” antibodies. Based on performance in immunocapture and other criteria, selected antibodies were subsequently tested for their ability to function as a probe to detect cell line EVs captured using anti-CD81 conjugated beads. (B, D, F) EVs from COV413A and SKMEL were captured with anti-CD81 bead conjugates and read out in the assay using monoclonal anti-FOLR1 (B), -BST2 (D), or -MUC1 (F) detection antibodies. Antibodies demonstrating a high degree of functionality and specificity were subject to additional testing and optimization (not shown).

**Supplemental Figure 4:**
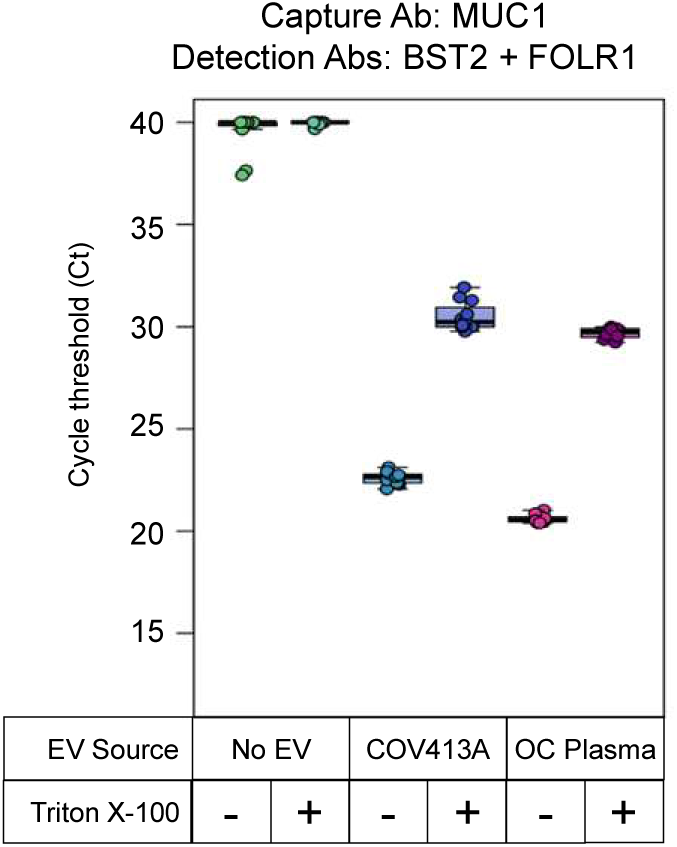
Detergent Sensitivity of Assay Performance. Buffer alone (No EV), EVs from COV413A or pooled human plasma from late-stage HGSOC patients’ plasma (OC Plasma) were assessed using MUC1 capture/FOLR1 + BST2 detection antibody combination as indicated. After capture on MUC1 antibody-bead conjugates, bead-bound EVs were incubated in buffer alone (-) or buffer containing 0.075% Triton-×100 (+) to disrupt the EV membranes as described in the methods. After a wash step to remove detergent, intact EVs were detected with anti-BST2 and anti-FOLR1 antibodies. Note that the presence of detergent results in a significant reduction in assay signal suggesting intact EVs are required for productive readout.

**Supplemental Figure 5:**
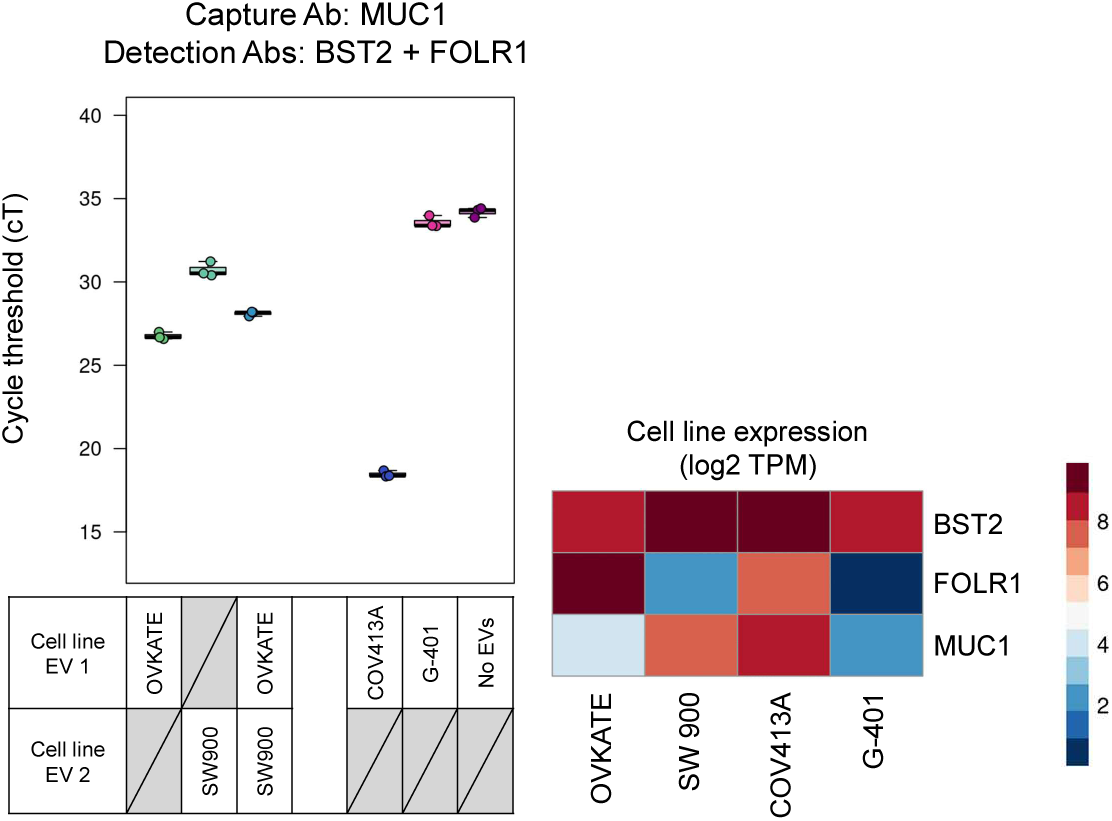
Demonstration of Single-EV Colocalization. EVs from OVKATE (BST2^high^, FOLR^high^, MUC1^low^), SW900 (BST2^high^, FOLR1^low^, MUC1^high^), COV413A (BST2^high^, FOLR1^high^, MUC1^high^), and G401 (BST2^high^, FOLR1^low^, MUC1^low^) were assayed using the BST2 capture/FOLR1 + MUC1 detection combination (A). EVs from OVKATE and SW900 were assayed independently or mixed 1:1 to determine whether oligo-conjugated antibodies on two independent EVs can “bridge” to generate a PCR signal.

**Supplemental Figure 6:**
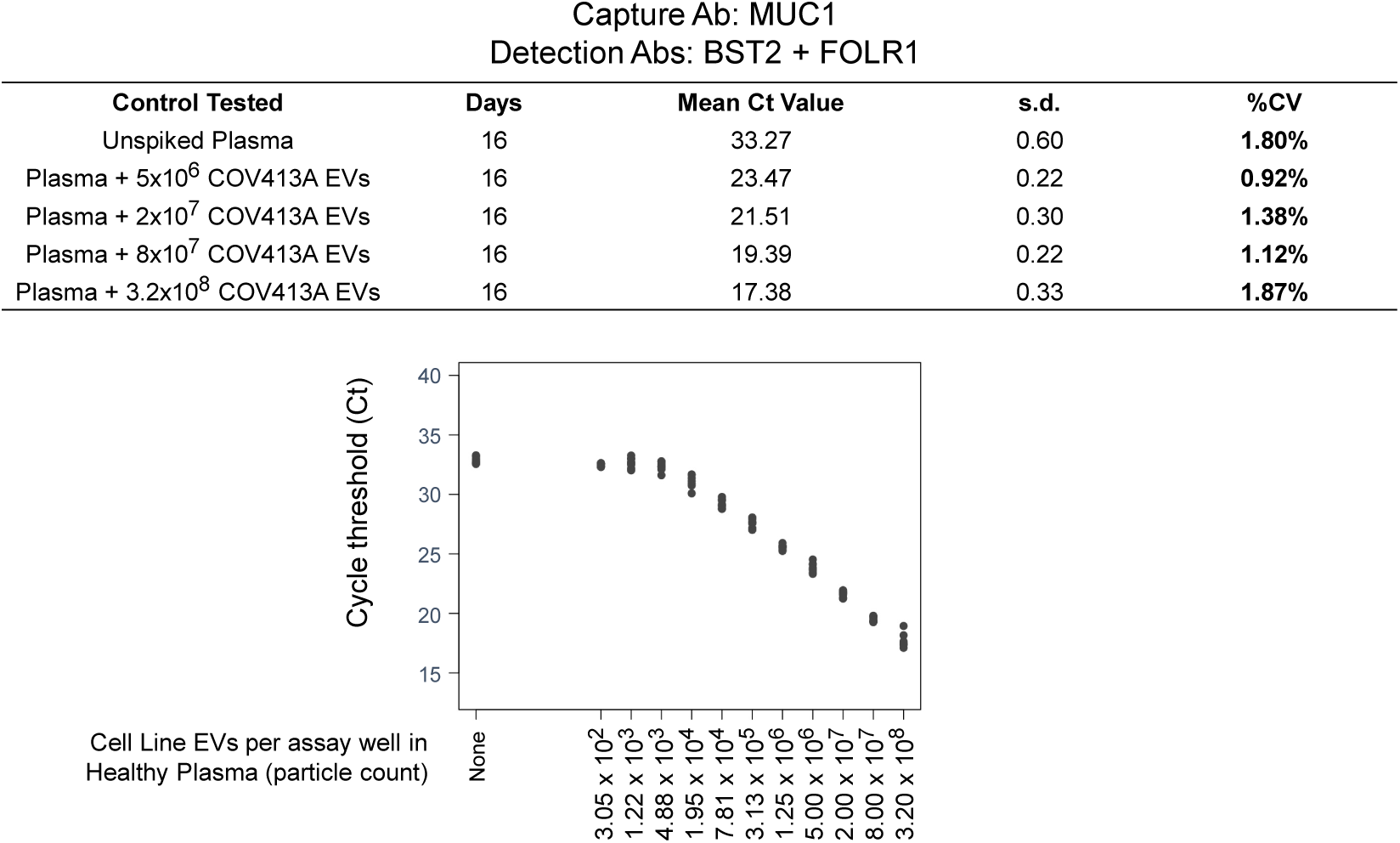
Reproducibility and Linearity of the Assay in Cell Line EV-Spiked Human Plasma. Representative reproducibility (top) and linearity (bottom) of assay performance in COV413A EV spiked human plasma using BST2/FOLR1/MUC1. Mean Ct values represent the mean of duplicate assays evaluated over 16 days. Points for each spike-in represent n=8 technical replicates. The last dilution point in linear range was 1.95 x 10^4^ which indicates a linear range over more than 15 Ct.

**Supplemental Figure 7.**
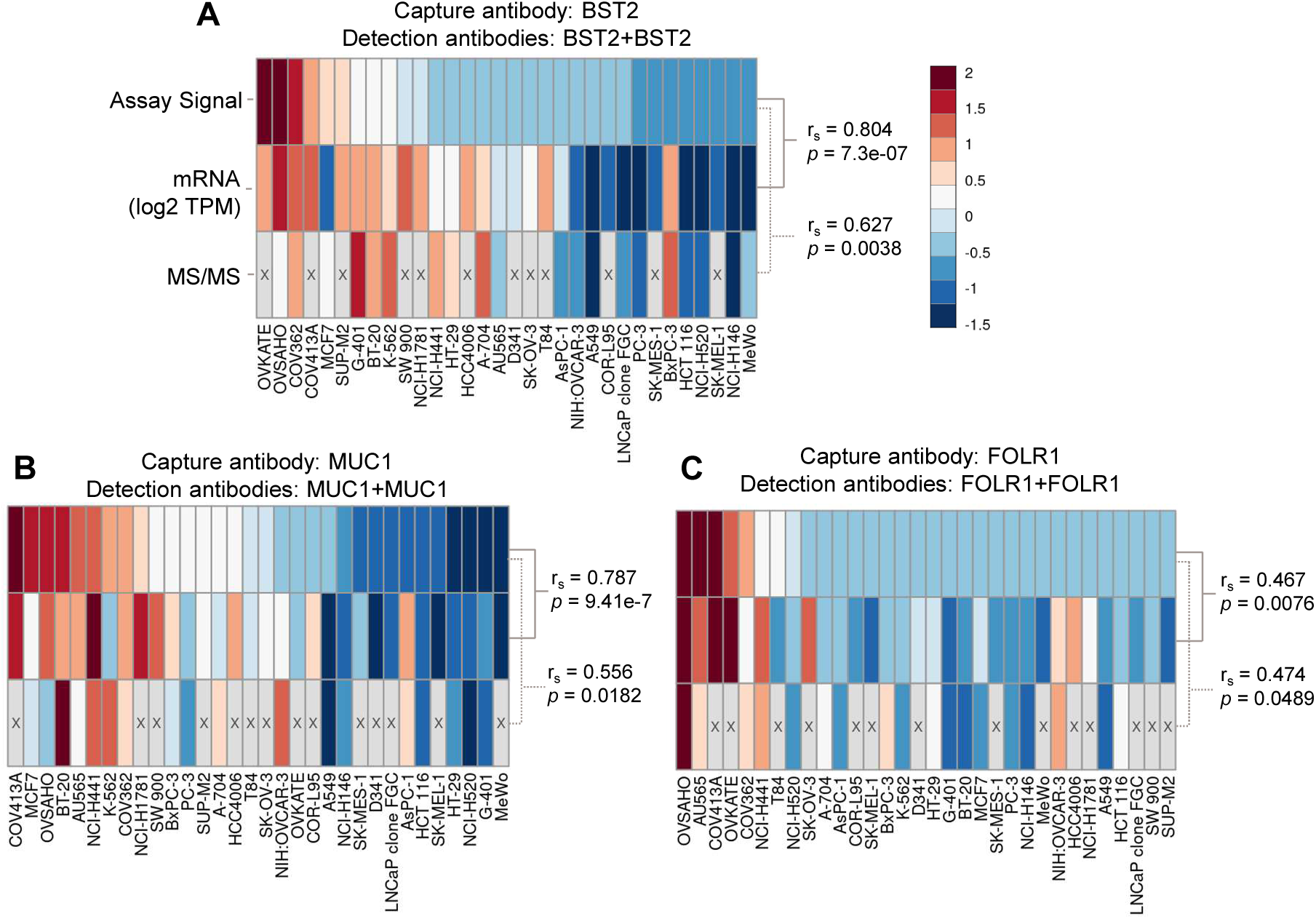
Correlation of assay signal with biomarker expression in cell line EVs. Heat maps comparing assay signal (top row), mRNA expression (middle row, log2 TPM), and protein abundance as determined by mass spectroscopy (bottom row, MS/MS) for the biomarkers tested in this study, BST2 (A), FOLR1 (B), and MUC1 (C). Gene expression and MS/MS data was sourced from the Cancer Cell Line Encyclopedia (CCLE) from the Broad Institute. The top row (Assay Signal) of each heat map is 40 minus the cycle threshold, the middle row (mRNA) is log2 transcripts per million (TPM), and the bottom row (MS/MS) is the log2 ratio of tandem mass tag (TMT) reporters vs control. All data was row scaled to allow comparison across data types. Spearman’s rank correlation (r_s_) and p-value (*p*) for each data set are shown.

**Supplemental Table 1:** Human cancer cell lines used in this study.

| Cell line name | Vendor | Catalog Number | RRID |
| --- | --- | --- | --- |
| A549 | ATCC | CCL-185 | CVCL_0023 |
| A704 | ATCC | HTB-45 | CVCL_1065 |
| AsPC1 | ATCC | CRL-1682 | CVCL_0152 |
| AU565 | ATCC | CRL-2351 | CVCL_1074 |
| BT-20 | ATCC | HTB-19 | CVCL_0178 |
| BxPC3 | ATCC | CRL-1687 | CVCL_0186 |
| COR-L-95 | Sigma-Aldrich/ECACC | 96020733 | CVCL_2418 |
| COV362 | Sigma-Aldrich/ECACC | 07071910 | CVCL_2420 |
| COV413A | Sigma-Aldrich/ECACC | 07071905 | CVCL_2422 |
| D341 | ATCC | HTB-187 | CVCL_0018 |
| G-401 | ATCC | CRL-1441 | CVCL_0270 |
| HCC4006 | ATCC | CRL-2871 | CVCL_1269 |
| HCT-116 | ATCC | CCL-247 | CVCL_0291 |
| HT-29 | ATCC | HTB-38 | CVCL_0320 |
| K-562 | ATCC | CCL-243 | CVCL_0004 |
| LnCaP (clone FGC) | ATCC | CRL-1740 | CVCL_1379 |
| MCF7 | ATCC | HTB-22 | CVCL_0031 |
| MeWo | ATCC | HTB-65 | CVCL_0445 |
| NCI-H146 | ATCC | HTB-173 | CVCL_1473 |
| NCI-H1781 | ATCC | CRL-5894 | CVCL_1494 |
| NCI-H441 | ATCC | HTB-174 | CVCL_1561 |
| NCI-H520 | ATCC | HTB-182 | CVCL_1566 |
| OVCAR3 | ATCC | HTB-161 | CVCL_0465 |
| OVISE | JCRB | JCRB1043 | CVCL_3116 |
| OVKATE | JCRB | JCRB1044 | CVCL_3110 |
| OVSAHO | JCRB | JCRB1046 | CVCL_3114 |
| PC-3 | ATCC | CRL-1345 | CVCL_0035 |
| SK-MEL-1 | ATCC | HTB-67 | CVCL_0068 |
| SK-MES-1 | ATCC | HTB-58 | CVCL_0630 |
| SK-OV-3 | ATCC | HTB-77 | CVCL_0532 |
| SUP-M2 | DSMZ | ACC 509 | CVCL_2209 |
| SW900 | ATCC | HTB-59 | CVCL_1731 |
| T47D | ATCC | HTB-133 | CVCL_0553 |
| T84 | ATCC | CCL-248 | CVCL_0555 |

Vendor Abbreviations: ATCC, American Type Culture Collection; ECACC, European Collection of Authenticated Cell Cultures; JCRB, Japanese Collection of Research Bioresources; DSMZ, Deutsche Sammlung von Mikroorganismen und Zellkulturen (German Collection of Microorganisms and Cell Cultures).

**Supplemental Table 2:** Antibodies used for western blot in this study.

| Target | Antibody Clone | Catalog # | Vendor | Dilution | RRID |
| --- | --- | --- | --- | --- | --- |
| CD63 | E1W3T | 52090 | CST | 1:1000 | AB_2924771 |
| CD81 | D3N2D | 56039 | CST | 1:1000 | AB_2924772 |
| CD9 | D801A | 13174 | CST | 1:1000 | AB_2798139 |
| Flotillin | D2V7J | 26836 | CST | 1:1000 | AB_2773040 |
| BST2 | D5V5Z | 19277 | CST | 1:1000 | AB_2798815 |
| FOLR1 | EPR20277 | ab235140 | Abcam | 1:1000 | AB_2924727 |
| TOM20 | D8T4N | 42406 | CST | 1:1000 | AB_2687663 |
| RPS3 | D50G7 | 9538 | CST | 1:1000 | AB_10622028 |
| CD81 | E2K9V | 52892 | CST | 1:1000 | AB_2924773 |
| GAPDH | D16H11 | 5174 | CST | 1:1000 | AB_10622025 |
| MUC 1-C | D5K9I | 16564 | CST | 1:1000 | AB_2798765 |
| Anti-Rabbit IgG, HRP-linked | Polyclonal | 7074 | CST | 1:1000 | AB_2099233 |
| Anti-Mouse IgG, HRP-linked | Polyclonal | 7076 | CST | 1:1000 | AB_330924 |
| Anti-Biotin IgG, HRP-linked | Polyclonal | 7075 | CST | 1:1000 | AB_10696897 |

Vendor Abbreviations: CST, Cell Signaling Technology

**Supplemental Table 3:** Antibodies used for capture and/or detection in this study.

| Name | Antibody Clone | Vendor | Catalog # | RRID |
| --- | --- | --- | --- | --- |
| Recombinant Anti-BST2/Tetherin antibody | EPR20202-169 | Abcam | ab243563 | AB_2941823 |
| Anti-BST2/Tetherin antibody [EPR20202-150] - BSA and Azide free | EPR20202-150 | Abcam | ab243561 | AB_2941851 |
| Anti-human CD317 | Y129 (HM1.24) | BD Biosciences | Custom order | Custom |
| BST2 Antibody (4F6) | 4F6 | Novus Biologicals | NBP2-29622 | AB_2941840 |
| BST2 monoclonal antibody (3H4) | 3H4 | Invitrogen | H00000684-M02 | AB_1136901 |
| Purified anti-human CD317 (BST2, Tetherin) Antibody | RS38E | Bio Legend | 348402 | AB_10588013 |
| CD317 (BST2, PDCA-1) Monoclonal Antibody (26F8), Functional Grade, eBioscience™ | 26F8 | Invitrogen Antibodies | 16-3179-82 | AB_1518775 |
| Anti-BST2/Tetherin antibody [EPR20202-169] - BSA and Azide free | EPR20202-169 | Abcam | ab243563 | AB_2941823 |
| Recombinant Anti-BST2/Tetherin antibody [EPR23597-202] - BSA and Azide free | EPR23597-202 | Abcam | ab272175 | AB_2941830 |
| Anti-BST2/Tetherin antibody - BSA and Azide free (Capture) | EPR20202-150 | Abcam | ab242967 | AB_2941821 |
| Anti-BST2/Tetherin antibody - BSA and Azide free (Detector) | EPR20202-169 | Abcam | ab243001 | AB_2941822 |
| Folate Receptor alpha monoclonal antibody (548908) | 548908 | ThermoFisher | MA5-23917 | AB_2609390 |
| Anti-Folate Binding Protein/FPB antibody [EPR23387-276], BSA and Azide free | EPR23387-276 | Abcam | ab273159 | AB_2941825 |
| Recombinant Anti-Folate Binding Protein/FPB antibody (Detector) [EPR23387-239] | EPR23387-239 | Abcam | ab278006 | AB_2941829 |
| Recombinant Anti-Folate Binding Protein/FPB antibody (Capture) [EPR23387-232] | EPR23387-232 | Abcam | ab278005 | AB_2941828 |
| Human FOLR1 Antibody | 548908 | R&D Systems | MAB5646 | AB_2278620 |
| FOLR1 Antibody (2G5C12) | 2G5C12 | Novus Biologicals | NBP2-61773 | AB_2941842 |
| Anti-Folate Binding Protein / FBP antibody [EPR20277] - BSA and Azide free | EPR20277 | Abcam | ab235140 | AB_2924727 |
| Anti-Human folate receptor 1 Recombinant Antibody (Farletuzumab) | Farletuzumab | Creative Biolabs | TAB-113 | AB_2941836 |
| Monoclonal Mouse anti Human FOLR1 / Folate Receptor Alpha Antibody | 3G12B7 | LSBio | LS-C682639 | AB_2941839 |
| Anti-Folate Binding Protein / FBP antibody [LK26] | LK26 | Abcam | ab3361 | AB_303740 |
| FOLR1 antibody | 2B4B7 | Proteintech | 60307-1-Ig | AB_2881421 |
| Human MUC-1 Antibody | 604804 | R&D Systems | MAB6298 | AB_10640403 |
| Anti-MUC1 antibody [EP1024Y] - Low endotoxin, Azide free | EP1024Y | Abcam | ab218998 | AB_2941819 |
| MUC1 (D9O8K) XP® Rabbit mAb | D9O8K | Cell Signaling Technology | 14161 | AB_2798408 |
| MUC1 (VU4H5) Mouse mAb | VU4H5 | Cell Signaling Technology | 4538 | AB_2148549 |
| Mouse anti Human CD227 | VU-3C6 | Bio-Rad | 6378-0135 | AB_620067 |
| MUC1 Monoclonal Antibody (GP1.4) | GP1.4 | Invitrogen Antibodies | MA1-06503 | AB_558126 |
| MUC1 Monoclonal Antibody (GP1.4) | GP1.4 | Novus Biologicals | NBP2-33174 | AB_2941841 |
| Epithelial Membrane Antigen (Concentrate) | E29 | DAKO | M061301-2 (1 mL) | AB_2148557 |
| Anti-Human MUC1 Recombinant Antibody (1B2) | 1B2 | Creative BioLabs | TAB-439MZ | AB_2941838 |
| Anti-MUC1 [12D10] | 12D10 | Absolute Antibody | Ab02843-1.1 | AB_2941833 |
| Human MUC-1 Antibody | 2336A | R&D Systems | MAB62981 | AB_2941844 |
| CD227 antibody \| C595 (NCRC48) | C595 (NCRC48) | Bio-Rad | MCA1742 | AB_322211 |
| MUC1 Antibody (SM3) | SM3 | Novus Biologicals | NB120-22711 | AB_789083 |
| MUC1 monoclonal antibody (115D8) | 115D8 | Invitrogen | MA1-35039 | AB_1070963 |
| Anti-MUC1 Antibody [115D8] - BSA and Azide free | 115D8 | Antibodies.com | A252591 | AB_2941835 |
| Anti-Human MUC1 (CD227) Antibody, Clone 16A | 16A | Stem cell Technologies | 60155 | AB_2941845 |
| Purified anti-human CD227 (MUC-1) Antibody | 16A | Bio Legend | 355602 | AB_2561642 |
| Anti-Human MUC1 Therapeutic Antibody (5E5) | 5E5 | Creative Biolabs | TAB-418MZ | AB_2941837 |
| Anti-MUC1 antibody [HMFG1 (aka 1.10.F3)] - BSA and Azide free | HMFG1 (aka 1.10.F3) | Abcam | ab230298 | AB_2941820 |
| Anti-MUC1 HMFG1 (1.10.F3) | HMFG1 | Absolute Antibody | AB02206-1.1 | AB_2941832 |
| Anti-MUC1 antibody [HMFG2] | HMFG2 | Abcam | ab245693 | AB_2941824 |
| Human MUC-1 Antibody | 604804 | R&D Systems | MAB6298 | AB_10640403 |

**Supplemental Table 4:** Patient Sample Demographics (Attached)

## References

Auber M, Svenningsen P. 2022. An estimate of extracellular vesicle secretion rates of human blood cells. J Extracell Biol 1. doi:10.1002/jex2.46

Avanzini S, Kurtz DM, Chabon JJ, Moding EJ, Hori SS, Gambhir SS, Alizadeh AA, Diehn M, Reiter JG. 2020. A mathematical model of ctDNA shedding predicts tumor detection size. Sci Adv 6. doi:10.1126/sciadv.abc4308

Bebelman MP, Smit MJ, Pegtel DM, Baglio SR. 2018. Biogenesis and function of extracellular vesicles in cancer. Pharmacol Ther 188:1–11.

Beer TM. 2021. Examining developments in multicancer early detection: highlights of new clinical data from recent conferences. Am J Manag Care 27:S347–S355.

Cabús L, Lagarde J, Curado J, Lizano E, Pérez-Boza J. 2022. Current challenges and best practices for cell-free long RNA biomarker discovery. Biomark Res 10:62.

Cizmar P, Yuana Y. 2017. Detection and Characterization of Extracellular Vesicles by Transmission and Cryo-Transmission Electron Microscopy. Methods Mol Biol 1660:221–232.

Crosby D, Bhatia S, Brindle KM, Coussens LM, Dive C, Emberton M, Esener S, Fitzgerald RC, Gambhir SS, Kuhn P, Rebbeck TR, Balasubramanian S. 2022. Early detection of cancer. Science 375:eaay9040.

Dang DK, Park BH. 2022. Circulating tumor DNA: current challenges for clinical utility. J Clin Invest 132. doi:10.1172/JCI154941

Desai A, Lovly CM. 2023. Challenges in the implementation of ultrasensitive liquid biopsy approaches in precision oncology. J Immunother Cancer 11:e006793.

Doyle LM, Wang MZ. 2019. Overview of Extracellular Vesicles, Their Origin, Composition, Purpose, and Methods for Exosome Isolation and Analysis. Cells 8. doi:10.3390/cells8070727

El Bairi K, Al Jarroudi O, Afqir S. 2021. Revisiting antibody-drug conjugates and their predictive biomarkers in platinum-resistant ovarian cancer. Semin Cancer Biol 77:42–55.

Eren E, Leoutsakos J-M, Troncoso J, Lyketsos CG, Oh ES, Kapogiannis D. 2022. Neuronal-Derived EV Biomarkers Track Cognitive Decline in Alzheimer’s Disease. Cells 11. doi:10.3390/cells11030436

Ferguson S, Weissleder R. 2020. Modeling EV Kinetics for Use in Early Cancer Detection. Adv Biosyst 4:e1900305.

Ferguson S, Yang KS, Weissleder R. 2022. Single extracellular vesicle analysis for early cancer detection. Trends Mol Med 28:681–692.

Gautam SK, Khan P, Natarajan G, Atri P, Aithal A, Ganti AK, Batra SK, Nasser MW, Jain M. 2023. Mucins as Potential Biomarkers for Early Detection of Cancer. Cancers 15. doi:10.3390/cancers15061640

Ghodasara A, Raza A, Wolfram J, Salomon C, Popat A. 2023. Clinical Translation of Extracellular Vesicles. Adv Healthc Mater e2301010.

Giampaolino P, Della Corte L, Foreste V, Vitale SG, Chiofalo B, Cianci S, Zullo F, Bifulco G. 2019. Unraveling a difficult diagnosis: the tricks for early recognition of ovarian cancer. Minerva Med 110:279–291.

Giampaolino P, Foreste V, Della Corte L, Di Filippo C, Iorio G, Bifulco G. 2020. Role of biomarkers for early detection of ovarian cancer recurrence. Gland Surg 9:1102–1111.

Han H, Jiang X. 2014. Overcome support vector machine diagnosis overfitting. Cancer Inform 13:145–158.

Ignatiadis M, Sledge GW, Jeffrey SS. 2021. Liquid biopsy enters the clinic - implementation issues and future challenges. Nat Rev Clin Oncol 18:297–312.

Januchowski R, Sterzyńska K, Zawierucha P, Ruciński M, Świerczewska M, Partyka M, Bednarek-Rajewska K, Brązert M, Nowicki M, Zabel M, Klejewski A. 2017. Microarray-based detection and expression analysis of new genes associated with drug resistance in ovarian cancer cell lines. Oncotarget 8:49944–49958.

Jerabkova-Roda K, Dupas A, Osmani N, Hyenne V, Goetz JG. 2022. Circulating extracellular vesicles and tumor cells: sticky partners in metastasis. Trends Cancer Res 8:799–805.

Johnsen KB, Gudbergsson JM, Andresen TL, Simonsen JB. 2019. What is the blood concentration of extracellular vesicles? Implications for the use of extracellular vesicles as blood-borne biomarkers of cancer. Biochim Biophys Acta Rev Cancer 1871:109–116.

Koshiyama M, Matsumura N, Konishi I. 2017. Subtypes of Ovarian Cancer and Ovarian Cancer Screening. Diagnostics (Basel) 7. doi:10.3390/diagnostics7010012

Kufe DW. 2009. Mucins in cancer: function, prognosis and therapy. Nat Rev Cancer 9:874–885.

Larssen P, Wik L, Czarnewski P, Eldh M, Löf L, Ronquist KG, Dubois L, Freyhult E, Gallant CJ, Oelrich J, Larsson A, Ronquist G, Villablanca EJ, Landegren U, Gabrielsson S, Kamali-Moghaddam M. 2017. Tracing Cellular Origin of Human Exosomes Using Multiplex Proximity Extension Assays. Mol Cell Proteomics 16:502–511.

Li Yuchen, He X, Li Q, Lai H, Zhang H, Hu Z, Li Yan, Huang S. 2020. EV-origin: Enumerating the tissue-cellular origin of circulating extracellular vesicles using exLR profile. Comput Struct Biotechnol J 18:2851–2859.

Liu J, Chen Y, Pei F, Zeng C, Yao Y, Liao W, Zhao Z. 2021. Extracellular Vesicles in Liquid Biopsies: Potential for Disease Diagnosis. Biomed Res Int 2021:6611244.

Lobb RJ, Becker M, Wen SW, Wong CSF, Wiegmans AP, Leimgruber A, Möller A. 2015. Optimized exosome isolation protocol for cell culture supernatant and human plasma. J Extracell Vesicles 4:27031.

Lone SN, Nisar S, Masoodi T, Singh M, Rizwan A, Hashem S, El-Rifai W, Bedognetti D, Batra SK, Haris M, Bhat AA, Macha MA. 2022. Liquid biopsy: a step closer to transform diagnosis, prognosis and future of cancer treatments. Mol Cancer 21:79.

Lundberg M, Eriksson A, Tran B, Assarsson E, Fredriksson S. 2011. Homogeneous antibody-based proximity extension assays provide sensitive and specific detection of low-abundant proteins in human blood. Nucleic Acids Res 39:e102.

Mariotto AB, Enewold L, Zhao J, Zeruto CA, Yabroff KR. 2020. Medical Care Costs Associated with Cancer Survivorship in the United States. Cancer Epidemiol Biomarkers Prev 29:1304–1312.

McGarvey N, Gitlin M, Fadli E, Chung KC. 2022. Increased healthcare costs by later stage cancer diagnosis. BMC Health Serv Res 22:1155.

Menon U, Gentry-Maharaj A, Burnell M, Singh N, Ryan A, Karpinskyj C, Carlino G, Taylor J, Massingham SK, Raikou M, Kalsi JK, Woolas R, Manchanda R, Arora R, Casey L, Dawnay A, Dobbs S, Leeson S, Mould T, Seif MW, Sharma A, Williamson K, Liu Y, Fallowfield L, McGuire AJ, Campbell S, Skates SJ, Jacobs IJ, Parmar M. 2021. Ovarian cancer population screening and mortality after long-term follow-up in the UK Collaborative Trial of Ovarian Cancer Screening (UKCTOCS): a randomised controlled trial. Lancet 397:2182–2193.

Menon U, Gentry-Maharaj A, Hallett R, Ryan A, Burnell M, Sharma A, Lewis S, Davies S, Philpott S, Lopes A, Godfrey K, Oram D, Herod J, Williamson K, Seif MW, Scott I, Mould T, Woolas R, Murdoch J, Dobbs S, Amso NN, Leeson S, Cruickshank D, McGuire A, Campbell S, Fallowfield L, Singh N, Dawnay A, Skates SJ, Parmar M, Jacobs I. 2009. Sensitivity and specificity of multimodal and ultrasound screening for ovarian cancer, and stage distribution of detected cancers: results of the prevalence screen of the UK Collaborative Trial of Ovarian Cancer Screening (UKCTOCS). Lancet Oncol 10:327–340.

Niemeyer CM, Adler M, Wacker R. 2005. Immuno-PCR: high sensitivity detection of proteins by nucleic acid amplification. Trends Biotechnol 23:208–216.

Osteikoetxea X, Sódar B, Németh A, Szabó-Taylor K, Pálóczi K, Vukman KV, Tamási V, Balogh A, Kittel Á, Pállinger É, Buzás EI. 2015. Differential detergent sensitivity of extracellular vesicle subpopulations. Org Biomol Chem 13:9775–9782.

Pavlik EJ, van Nagell JR Jr. 2013. Early detection of ovarian tumors using ultrasound. Womens Health 9:39–55; quiz 56–7.

Pinsky PF. 2014. Assessing the benefits and harms of low-dose computed tomography screening for lung cancer. Lung Cancer Manag 3:491–498.

Siegel RL, Miller KD, Wagle NS, Jemal A. 2023. Cancer statistics, 2023. CA Cancer J Clin 73:17– 48.

Stiller C, Viktorsson K, Paz Gomero E, Hååg P, Arapi V, Kaminskyy VO, Kamali C, De Petris L, Ekman S, Lewensohn R, Karlström AE. 2021. Detection of Tumor-Associated Membrane Receptors on Extracellular Vesicles from Non-Small Cell Lung Cancer Patients via Immuno-PCR. Cancers 13. doi:10.3390/cancers13040922

Sung H, Ferlay J, Siegel RL, Laversanne M, Soerjomataram I, Jemal A, Bray F. 2021. Global Cancer Statistics 2020: GLOBOCAN Estimates of Incidence and Mortality Worldwide for 36 Cancers in 185 Countries. CA Cancer J Clin 71:209–249.

Tavoosidana G, Ronquist G, Darmanis S, Yan J, Carlsson L, Wu D, Conze T, Ek P, Semjonow A, Eltze E, Larsson A, Landegren UD, Kamali-Moghaddam M. 2011. Multiple recognition assay reveals prostasomes as promising plasma biomarkers for prostate cancer. Proc Natl Acad Sci U S A 108:8809–8814.

Théry C, Witwer KW, Aikawa E, Alcaraz MJ, Anderson JD, Andriantsitohaina R, Antoniou A, Arab T, Archer F. 2018. Minimal information for studies of extracellular vesicles 2018 (MISEV2018): a position statement of the International Society for Extracellular Vesicles and update of the MISEV2014 guidelines. J Extracell Vesicles 7:1535750.

Trinidad C, Pathak H, Cheng S, Tzeng S-C, Madan R, Sardiu M, Bantis L, Deighan C, Jewell A, Zeng Y, Godwin A. 2023. Lineage specific extracellular vesicle-associated protein biomarkers for the early detection of high grade serous ovarian cancer. Res Sq. doi:10.21203/rs.3.rs-2814022/v1

US Preventive Services Task Force. 2009. Screening for breast cancer: U.S. Preventive Services Task Force recommendation statement. Ann Intern Med 151:716–26, W-236.

US Preventive Services Task Force, Davidson KW, Barry MJ, Mangione CM, Cabana M, Caughey AB, Davis EM, Donahue KE, Doubeni CA, Krist AH, Kubik M, Li L, Ogedegbe G, Owens DK, Pbert L, Silverstein M, Stevermer J, Tseng C-W, Wong JB. 2021a. Screening for Colorectal Cancer: US Preventive Services Task Force Recommendation Statement. JAMA 325:1965–1977.

US Preventive Services Task Force, Grossman DC, Curry SJ, Owens DK, Barry MJ, Davidson KW, Doubeni CA, Epling JW Jr, Kemper AR, Krist AH, Kurth AE, Landefeld CS, Mangione CM, Phipps MG, Silverstein M, Simon MA, Tseng C-W. 2018. Screening for Ovarian Cancer: US Preventive Services Task Force Recommendation Statement. JAMA 319:588–594.

US Preventive Services Task Force, Krist AH, Davidson KW, Mangione CM, Barry MJ, Cabana M, Caughey AB, Davis EM, Donahue KE, Doubeni CA, Kubik M, Landefeld CS, Li L, Ogedegbe G, Owens DK, Pbert L, Silverstein M, Stevermer J, Tseng C-W, Wong JB. 2021b. Screening for Lung Cancer: US Preventive Services Task Force Recommendation Statement. JAMA 325:962–970.

van Nagell JR Jr, DePriest PD, Ueland FR, DeSimone CP, Cooper AL, McDonald JM, Pavlik EJ, Kryscio RJ. 2007. Ovarian cancer screening with annual transvaginal sonography: findings of 25,000 women screened. Cancer 109:1887–1896.

van Niel G, D’Angelo G, Raposo G. 2018. Shedding light on the cell biology of extracellular vesicles. Nat Rev Mol Cell Biol 19:213–228.

Varaganti P, Buddolla V, Lakshmi BA, Kim Y-J. 2023. Recent advances in using folate receptor 1 (FOLR1) for cancer diagnosis and treatment, with an emphasis on cancers that affect women. Life Sci 326:121802.

Veerman RE, Teeuwen L, Czarnewski P, Güclüler Akpinar G, Sandberg A, Cao X, Pernemalm M, Orre LM, Gabrielsson S, Eldh M. 2021. Molecular evaluation of five different isolation methods for extracellular vesicles reveals different clinical applicability and subcellular origin. J Extracell Vesicles 10:e12128.

Weibrecht I, Leuchowius K-J, Clausson C-M, Conze T, Jarvius M, Howell WM, Kamali-Moghaddam M, Söderberg O. 2010. Proximity ligation assays: a recent addition to the proteomics toolbox. Expert Rev Proteomics 7:401–409.

Willms E, Cabañas C, Mäger I, Wood MJA, Vader P. 2018. Extracellular Vesicle Heterogeneity: Subpopulations, Isolation Techniques, and Diverse Functions in Cancer Progression. Front Immunol 9:738.

Yang L-Q, Hu H-Y, Han Y, Tang Z-Y, Gao J, Zhou Q-Y, Liu Y-X, Chen H-S, Xu T-N, Ao L, Xu Y, Che X, Jiang Y-B, Xu C-W, Zhang X-C, Jiang Y-X, Heger M, Wang X-M, Cheng S-Q, Pan W-W. 2022. CpG-binding protein CFP1 promotes ovarian cancer cell proliferation by regulating BST2 transcription. Cancer Gene Ther 29:1895–1907.

Yu D, Li Y, Wang M, Gu J, Xu W, Cai H, Fang X, Zhang X. 2022. Exosomes as a new frontier of cancer liquid biopsy. Mol Cancer 21:56.

Yu W, Hurley J, Roberts D, Chakrabortty SK, Enderle D, Noerholm M, Breakefield XO, Skog JK. 2021. Exosome-based liquid biopsies in cancer: opportunities and challenges. Ann Oncol 32:466–477.

